# Lower Soil Carbon Loss Due to Persistent Microbial Adaptation to Climate Warming

**DOI:** 10.1101/2020.02.23.961300

**Authors:** Xue Guo, Qun Gao, Mengting Yuan, Gangsheng Wang, Xishu Zhou, Jiajie Feng, Zhou Shi, Lauren Hale, Linwei Wu, Aifen Zhou, Renmao Tian, Feifei Liu, Bo Wu, Lijun Chen, Chang Gyo Jung, Shuli Niu, Dejun Li, Xia Xu, Lifen Jiang, Arthur Escalas, Liyou Wu, Zhili He, Joy D. Van Nostrand, Daliang Ning, Xueduan Liu, Yunfeng Yang, Edward. A.G. Schuur, Konstantinos T. Konstantinidis, James R. Cole, C. Ryan Penton, Yiqi Luo, James M. Tiedje, Jizhong Zhou

## Abstract

Soil microbial respiration is an important source of uncertainty in projecting future climate and carbon (C) cycle feedbacks. Despite intensive studies for two decades, the magnitude, direction, and duration of such feedbacks are uncertain, and their underlying microbial mechanisms are still poorly understood. Here we examined the responses of soil respiration and microbial community structure to long-term experimental warming in a temperate grassland ecosystem. Our results indicated that the temperature sensitivity of soil microbial respiration (i.e., *Q*_10_) persistently decreased by 12.0±3.7% across 7 years of warming. Integrated metagenomic and functional analyses showed that microbial community adaptation played critical roles in regulating respiratory acclimation. Incorporating microbial functional gene abundance data into a microbially-enabled ecosystem model significantly improved the modeling performance of soil microbial respiration by 5–19%, compared to the traditional non-microbial model. Model parametric uncertainty was also reduced by 55–71% when gene abundances were used. In addition, our modeling analyses suggested that decreased temperature sensitivity could lead to considerably less heterotrophic respiration (11.6±7.5%), and hence less soil C loss. If such microbially mediated dampening effects occur generally across different spatial and temporal scales, the potential positive feedback of soil microbial respiration in response to climate warming may be less than previously predicted.

## Introduction

Soil stores large quantities of organic carbon (C), about three times more C than the Earth’s atmosphere ^1,2^. Soil respiration is the largest single source of carbon dioxide (CO_2_) from terrestrial ecosystems to the atmosphere, whose magnitude is about ten times larger than anthropogenic emissions ^3^. Soil total respiration (*R*_t_) includes both autotrophic respiration (*R*_a_) from plant root growth and root biomass maintenance, and heterotrophic respiration (*R*_h_) from microbial decomposition of litter and soil organic matter (SOM). Various short-term experiments show that soil respiration increases exponentially with temperature ^4^, which has been used as a general relationship to parameterize ecosystem and Earth System Models (ESMs) ^5^. If the near-exponential short-term relationship of soil respiration and temperature holds for the long-term (years to decades), climate warming will trigger a sharp increase in ecosystem respiration. Such an increase could then result in a strong positive feedback to the global C cycle ^6^, which is dependent on the responses of *R*_h_ and the dynamics of detrital inputs under warming ^7^. Therefore, it is particularly important to accurately evaluate soil *R*_h_ and its response to climate warming. However, partitioning *R*_t_ into *R*_a_ and *R*_h_ is one of the main challenges in both experiment- and model-based global change research ^8^. Consequently, soil respiration is a poorly understood key C flux in the global C cycle and is an important source of the uncertainty in climate projections ^9–11^.

Microorganisms can dramatically adjust their respiratory responses to temperature over long terms (years) via changing their metabolism and community structure ^12^. Several climate change experiments demonstrated that soil respiration was stimulated in the short term, followed by a dampened effect of warming later ^13–15^. This phenomenon is referred to as respiratory acclimation. The existence of respiratory acclimation is of critical importance as the greater the global respiratory acclimation, the weaker the positive feedback between climate warming and ecosystem CO_2_ release ^16^. However, the existence and the degree of soil respiratory acclimation is extremely uncertain, especially in the field and over a long duration (years to decades) ^9,10,17^. Whether respiratory acclimation can persist over time is not clear. Moreover, the mechanisms controlling soil respiratory acclimation have been intensively debated ^4,14,17–19^, and include warming-induced substrate depletion ^17,19^ or evolutionary adaptation via changes in microbial community ^13,14^. These two mechanisms may lead to different soil C loss in a warmer world ^14,19^. While the former could lead to a depletion of labile C pools, releasing more C into the atmosphere through microbial respiration if more plant-derived C is available under warming, the latter could result in less soil labile C loss due to microbial community adaptation to the rising temperature (warming) ^14^. Therefore, knowledge about microbial respiratory acclimation and its underlying mechanisms will be central to making better predictions of terrestrial C cycling feedbacks. However, one grand challenge in climate change biology is to integrate microbial community information, particularly omics information, into ecosystem models to improve their predictive ability for projecting future climate and environmental changes ^20^. More specifically, parameter values for various microbial processes are poorly constrained by experimental observations, which becomes one of the significant uncertainty sources leading to low confidence in carbon-climate feedback projections ^21^. Hence, using omics-enabled experimental observations to improve model parameter estimations could greatly help to refine the projected magnitude of the carbon-climate feedbacks.

Soil microbial communities are very complex in structure and are sensitive to changes in environmental conditions ^14^, so information obtained from a single time point provides only a snapshot of the microbial community, and is not suitable for ecosystem model simulation. To modeling microbial respiratory responses to climate warming, long-term experiments under more realistic field-settings with time-series microbial data are needed. Otherwise, it will be difficult to determine the direction, magnitude, and duration of biospheric feedbacks to climate change ^15,22^. Therefore, a new warming experiment site with sandy soil and dominance of C_3_ grasses was established in a native, tall-grass prairie ecosystem of the US Great Plains in Central Oklahoma (34° 59’ N, 97° 31’ W) in July 2009 ^23^. Soil samples archived every year right after the continuous warming by infrared radiators (+3 °C) were analyzed by integrated metagenomics technologies.

In this study, we examined the temperature responses of soil *R*_h_ (> 7 years) and their underlying mechanisms. Our main objectives were to answer the following questions: (i) How does long-term experimental warming affect the temperature responses of soil microbial respiration over time? (ii) Whether or not acclimation of microbial respiration occurs persistently across years under warming and by what underlying mechanisms? (iii) Can the microbial mechanisms underlying soil respiration be incorporated into ecosystem models to improve model performance and reduce model uncertainty? We hypothesize that soil microbial respiratory acclimation exists persistently over the long-term and that microbial community adaptation plays critical roles in regulating such respiratory acclimation. If true, incorporating metagenomics-based microbial functional information will significantly increase confidence in model simulations and therefore improve model predictions.

## Results and discussion

### Overall ecosystem changes under long-term warming

The plots in the warming experiment site have been subjected to continuous warming for over 7 years ^7^. On average, experimental warming significantly (p < 0.01) increased daily air temperature by 1.3 °C, and daily mean soil temperature at 7.5 cm by 2.8 °C (Fig. 1a). Experimental warming significantly (p < 0.01) decreased soil moisture by 6.4% (Fig. 1b). Consistent with previous reports ^14^, warming significantly (p = 0.01) shifted plant community structure. Specifically, C_3_ plant biomass was significantly (p < 0.01) lower under warming than control, but no significant change was observed in C_4_ and total plant biomass (Fig. S1a), which results in a plant community shift towards relatively more C_4_ plants. Although the statistical test is not significant, the gross primary production (GPP) was slightly increased by warming (Fig. 1c). Meanwhile, the net ecosystem exchange (NEE) was higher under warming than control due to lower ecosystem respiration (ER), suggesting that the whole ecosystem acted as a C sink under the climate warming scenario (Fig. 1c). In addition, no overall differences were detected in total organic C (TOC), total nitrogen (TN) and soil pH (Fig. S1b and c), but the amount of (NO_3_^−^) was significantly higher under warming than control (Fig. S1c). These alterations in ecosystem variables by warming are expected to lead to changes in soil respirations and microbial community functions.

**Fig. 1.**
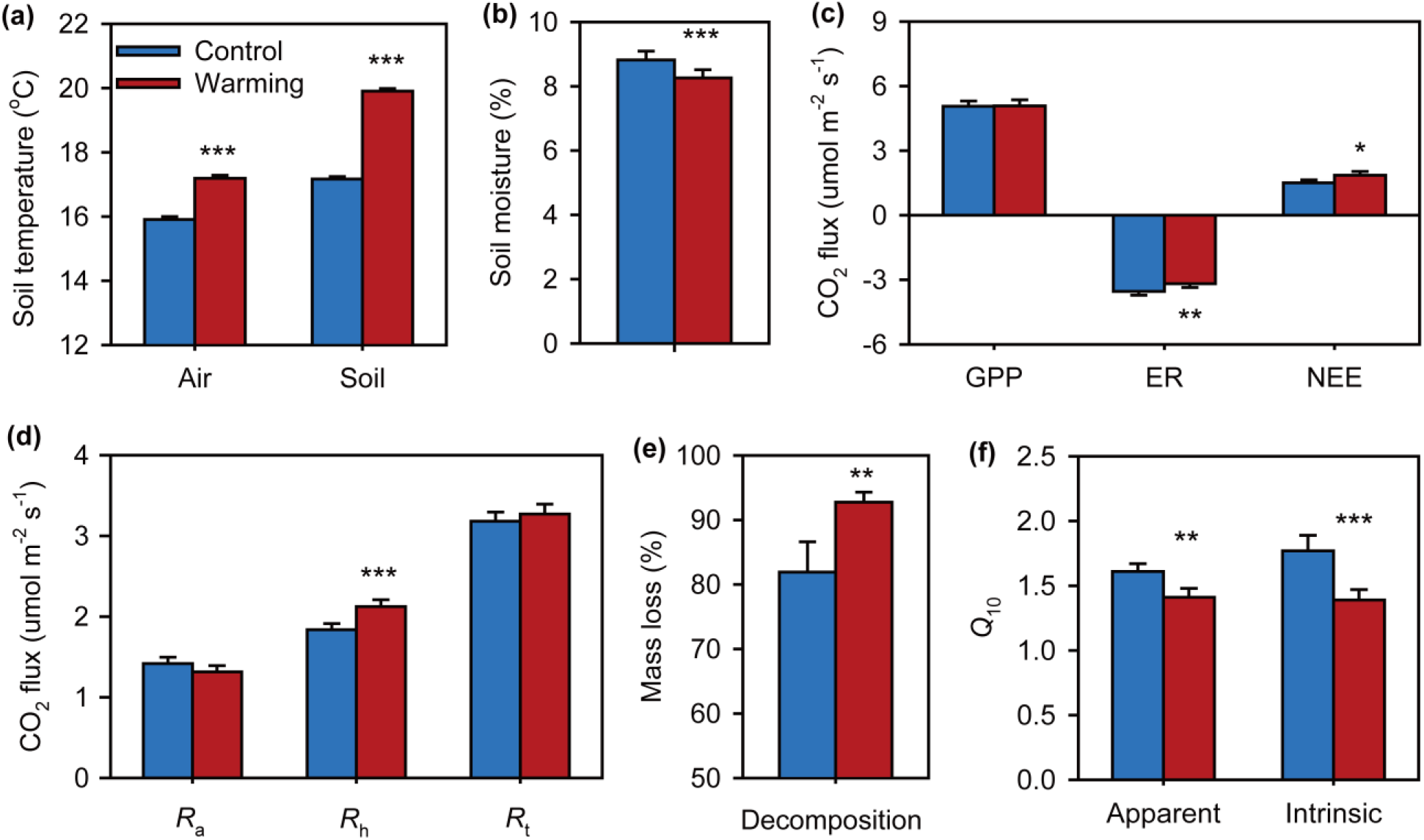
Warming effects on soil variables and ecosystem C fluxes. **(a)** Air and soil surface (7.5 cm) temperatures averaged from 2010 to 2016. (**b)** Soil moisture averaged from 2010 to 2016. (**c**) Ecosystem C fluxes, which were estimated on the basis of the C amount from CO_2_ emissions averaged from 2010 to 2016. GPP, gross primary productivity; ER, ecosystem respiration; NEE, net ecosystem C exchange. Positive values indicate C sink, and negative values represent C source. (**d**) *in situ* soil respirations averaged from 2010 to 2016. *R*_a_, autotrophic respiration; *R*_h_, heterotrophic respiration; *R*_t_, soil total respiration. (**e**) Decomposition rate of standard cellulose filter paper (mass loss) in the field determined in 2016. (**f**) Apparent and intrinsic temperature sensitivity (*Q*_10_) of heterotrophic respiration (*R*_*h*_) averaged from 2010 to 2016. Apparent *Q*_10_ is estimated by fitting the curve of *R*_*h*_ versus soil temperature based on the *Q*_10_ method. Intrinsic *Q*_10_ is derived by calibrating the MEND model. Error bars represent standard error of the mean. The differences between warming and control were tested by repeated measures ANOVA, indicated by *** when p < 0.01, ** when p < 0.05, * when p < 0.10.

### Temperature sensitivity of soil microbial respiration under warming

Soil surface CO_2_ efflux was measured by using shallow (2-3cm) PVC collars for *R*_t_ and deep (70cm) PVC tubes for *R*_h_, with the differences between *R*_t_ and *R*_h_ calculated as *R*_a_ (Fig. S2 and Methods). Warming significantly (p < 0.01) stimulated *R*_h_ by 8.0–28.1% across all years, which is consistent with results from a filter paper decomposition experiment that showed significantly (p<0.01) higher decomposition rates under warming (Fig. 1e). However, warming appeared to suppress *R*_a_, although it was not statistically significant (Fig. 1d), which may result from the decreased root activities along warming-induced plant community shift ^7^. More than half of *R*_t_ (58% and 65% for the control and warming plots) was from heterotrophic respiration, indicating that soil microbial community greatly contribute to soil CO_2_ efflux ^14^. No significant decline of *R*_h_/*R*_t_ ratio was observed in warmed and control plots through time, suggesting that soil C input in the form of plant litter may substantially contribute to the stability of soil C when plant roots were excluded. Due to the opposing responses of *R*_a_ and *R*_h_ to warming, *R*_t_ exhibited no significant change by warming across all years (Fig. 1d). Since our main interest is the response of microbial litter and SOM decomposition to warming, we primarily focused on *R*_h_ for the majority of the following analyses.

To examine the apparent temperature sensitivity (*Q*_10_) of microbial respiration, the measured field *R*_h_ data in each year were fitted to the *Q*_10_-based exponential equation ^4^ (see Methods). Significant (p < 0.05) or marginally significant (p < 0.10) apparent *Q*_10_ estimates were observed under both control and warming treatments in all years except 2011 (Table S1). In average, the apparent *Q*_10_ estimates were significantly (p = 0.03) higher under control (1.61±0.06) than warming (1.41± 0.07), suggesting a 12.0±3.7% decrease in the temperature sensitivity of soil *R*_h_ across 7 years of warming (Fig. 1f). However, the apparent temperature sensitivity estimate based on the field measurements are influenced by various other factors beyond temperature, including soil moisture, plants-derived substrate quality and availability, nutrient limitation influencing microbial enzyme production, experimental duration, and/or spatial heterogeneity, as well as uncertainty in instrumental measurements ^4,8^.

To further delineate the intrinsic temperature sensitivity of SOM decomposition, ecosystem model-based inverse analysis was performed to untangle various complex soil processes ^8,14,18^ using the Microbial-ENzyme Decomposition (MEND) model (Fig. S3a), which has been evaluated from laboratory to global scale ^24–26^. By fitting all 7-year respiration data together, the model-based intrinsic *Q*_10_ under warming was 1.39±0.09, significantly lower (p< 0.01) than that under control (1.77±0.12) (Fig. 1f). The intrinsic *Q*_10_ values from our model-data fusion approach were comparable with the measured apparent *Q*_10_ under both control and warming. Altogether, the above results indicate that there was a strong and persistent acclimation of heterotrophic respiration under warming over the last 7 years.

### Mechanisms of the persistent decrease in temperature sensitivity of microbial respiration

The persistent decrease in temperature sensitivity of soil microbial respiration across different years under warming could be due to substrate depletion under warming. It has been argued that soil labile C becomes depleted by increased respiration in response to warming, which leads to a subsequent reduction in the rate of soil respiration ^10^. In this study, several lines of evidence suggest that the decreased temperature sensitivity of microbial respiration was unlikely due to substrate depletion. First, GPP and NEE were similar or higher under warming than control (Fig. 1c), suggesting that soil C input as plant litter and root exudates should be similar or even higher under warming than control. Also, our BIOLOG results revealed that microbial metabolism underpinning the utilization ability of most labile substrates were considerably higher under warming than control (Fig. S4). The measured mean annual soil C from 2010 to 2016 remained unchanged (Fig. S1c), which do not support the expectation garnered from the substrate depletion hypothesis. These results suggested that the reduced temperature sensitivity of soil respiration appears to be less likely due to substrate depletion.

The adaptive changes in microbial community composition and functional structure could also lead to the reduced temperature sensitivity of microbial respiration. To test this hypothesis, soil microbial communities of individual samples from 2010 to 2016 were all analyzed with deep amplicon sequencing of the 16S rRNA gene for bacteria and archaea, and the ITS for fungi, metagenomic shotgun sequencing, and functional gene arrays (GeoChip 5.0; Table S2). Permutational multivariate analysis revealed that experimental warming significantly shifted microbial community taxonomic and functional structure (Table 1). These shifts were tightly linked to environmental factors, such as soil temperature, soil moisture, pH and climate conditions as revealed by the Mantel test (Fig. 2a and S5) and canonical correspondence analyses (CCA) (Fig. S6). Interestingly, considerably less unexplained community variations were obtained based on GeoChip data (59.2%) than 16S (73.0%), ITS (77.4%) and shotgun sequencing data (73.3%) (Fig. S7), indicating that GeoChip-based detection could be more effective to catch the community dynamics in response to the changes in plant diversity, soil conditions, and time. In addition, structural equation modeling (SEM)-based analysis indicated that soil temperature, moisture and drought index could strongly affect soil *R*_h_ by altering microbial functional diversity and structure (Fig. 2b).

**Table 1.** Significance tests of the effects of warming and time on microbial community structures with permutational multivariate analysis of variance. Permutational multivariate analysis of variance (Adonis) was used based on Bray-Curtis dissimilarity matrices. The two-way repeated-measures ANOVA model was set as “dissimilarity~warming×year+block” using function adonis in R package vegan. The degree of freedom was 1 for warming treatment, 6 for year and 39 for residuals. Significant effects (P ≤ 0.05) were shown in bold text. EcoFUN-MAP is a method designed for annotating metagenomic sequences by comparing them with functional genes used to fabricate GeoChip.

| Effects | 16S |  | ITS |  | GeoChip |  | Metagenomic sequencing |  | Metagenome based EcoFUN-MAP |  |
| --- | --- | --- | --- | --- | --- | --- | --- | --- | --- | --- |
|  | F | P | F | P | F | P | F | P | F | P |
| Warming (W) | 4.200 | **0.001** | 2.314 | **0.001** | 2.505 | **0.026** | 8.059 | **0.001** | 2.924 | **0.001** |
| Year (Y) | 2.432 | **0.001** | 1.595 | **0.001** | 12.216 | **0.001** | 4.398 | **0.001** | 2.323 | **0.001** |
| W × Y | 1.178 | 0.092 | 1.055 | 0.224 | 1.385 | 0.092 | 1.350 | 0.170 | 1.135 | 0.084 |

**Fig. 2.**
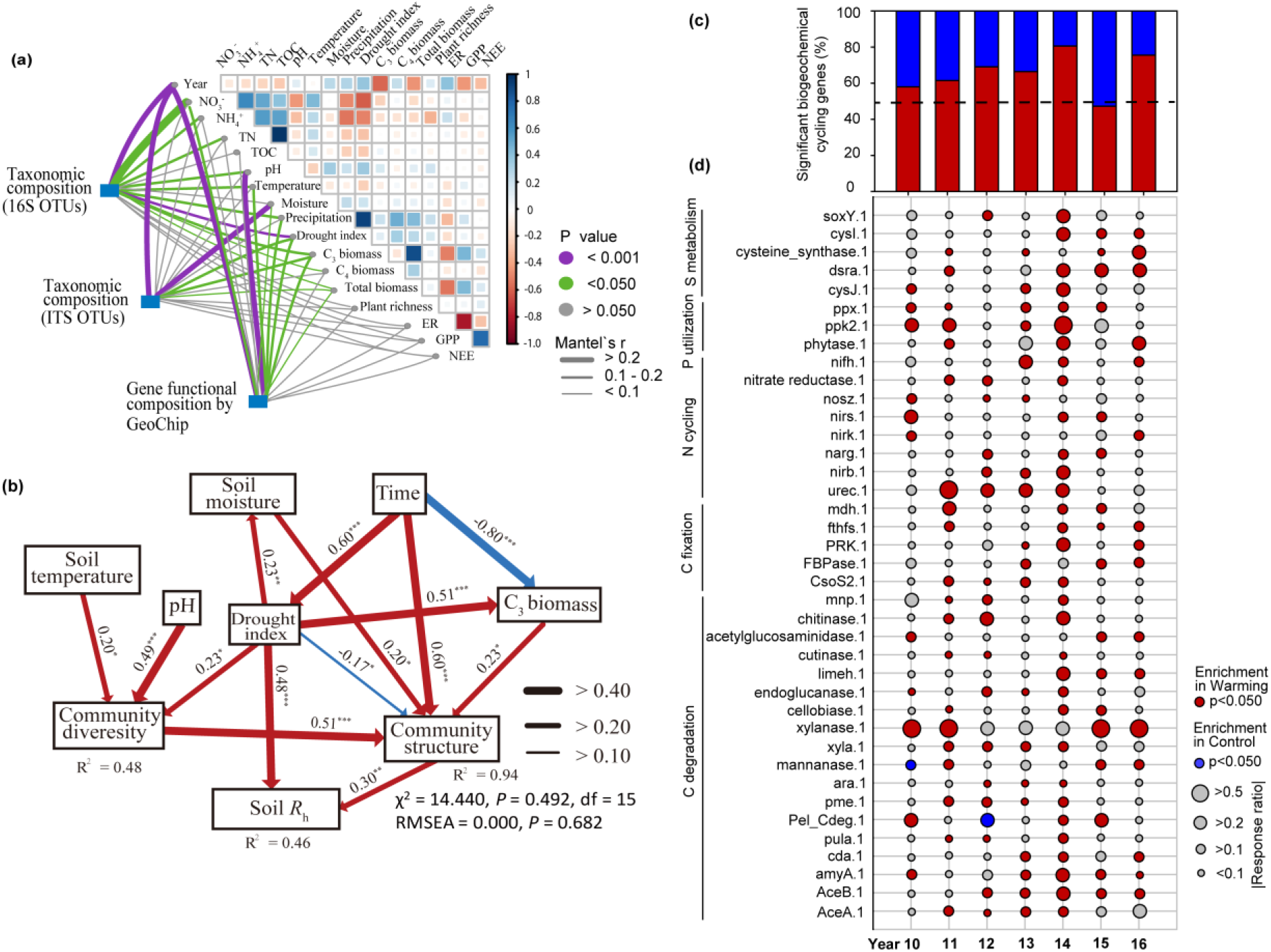
Feedback mechanisms of soil microbial communities to warming. (**a**) Pairwise comparisons of environmental factors with a color gradient denoting Pearson’s correlation coefficients. Taxonomic (based on 16S rRNA gene and ITS OTUs) and functional (based on GeoChip data) community structures were related to each environmental factor by Mantel tests. Edge width corresponds to the Mantel’s r statistic for the corresponding distance correlations, and edge color denotes the statistical significance. (**b**) The structural equation model (SEM) showing causal relationships among environmental factors, community diversity (Shannon index based on GeoChip) and structure (the first axis of NMDS analysis of GeoChip data), and heterotrophic respiration (*R*_h_). Red and blue arrows represent significant positive and negative pathways, respectively. Arrow width is proportional to the strength of the relationship and bold numbers represent the standard path coefficients, and the p values of path coefficients are indicated by *** when P < 0.001, ** when P < 0.01, * when P < 0.05. R^2^ indicates the proportion of the variance explained for each dependent variable in the model. (**c**) Biogeochemical cycling genes significantly changed by warming from 2010 to 2016 according to GeoChip data. Biogeochemical cycling genes included all genes involved in C degradation, C fixation, N cycling, phosphorus (P) utilization and sulfur (S) metabolism. Significance is based on response ratio of each gene with 95% confidence intervals of abundance differences between warmed and control treatments. Dash line represents that the abundance of warming-stimulated (red) genes are in good agreement with the abundance of warming-inhibited (blue) genes. (**d**) Bubble plot illustrating the enrichment of key biogeochemical cycling genes under warming (W) and control (C) treatments according to GeoChip data. Bubble color represents the significance (p-value) of gene enrichment based on response ratios. Bubble size represents the relative changes of gene enrichment based on response ratios. The biogeochemical cycling processes for these genes are shown in plot, and the full names of the genes in this plot are listed in Supplementary Table S5.

Warming-induced shifts of microbial functional diversity and structure led to significant changes of biogeochemical cycling processes, including C cycling (e.g., C degradation, C fixation) and nutrient-cycling processes (e.g., N fixation, denitrification, nitrification), phosphorus utilization and sulfur metabolism. Overall, the total abundance of biogeochemical cycling genes significantly (p < 0.05) stimulated by warming were considerably higher (58%~80%) than those significantly inhibited by warming (20%~42%) in all years except 2015 (Fig. 2c), although the interannual variations of environmental factors greatly influenced the composition of biogeochemical cycling genes. Similar pattern was also observed in microbial functional genes involved in C degradation (Fig. S8a), including those important for degrading starch (e.g., *amyA* encoding α-amylase), hemicellulose (e.g., *ara* encoding arabinofuranosidase), cellulose (e.g., cellobiase), chitin (e.g., chitinase) and vanillin/lignin (e.g., *mnp* encoding manganese peroxidase). More specifically, larger numbers of individual genes involved in degrading various soil organic carbon were significantly increased by warming (95% confidence interval; Fig. 2d and Fig. S9) in most of the years, despite that warming effects on these C-degrading genes substantially changed across different years. The significant enrichment of C-degrading genes under warming may potentially enhance soil C degradation. In addition, the total abundances of warming-stimulated genes involved in N cycling (e.g., N fixation, denitrification, and nitrification), phosphorus utilization, and sulfur metabolism were higher than those of warming-inhibited genes in most of the years (Fig. 2d and Fig. S8b-d), suggesting that the rates of nutrient-cycling processes could be stimulated by warming. Further analyses by CCA and Mantel test revealed that most of the genes important to C degradation and nutrient cycling had strong correlations to the *R*_h_, *R*_t_, and *Q*_10_ (Table S3 and S4), indicating that these functional genes are important in controlling the dynamics of soil respirations. In general, GeoChip hybridization data exhibited stronger correlations to various functional parameters than shotgun sequencing data, particularly for the heterotrophic *Q*_10_ (Table S3 and S4). All the above results indicated that the changes of microbial community composition and function are crucial for the reduced temperature sensitivity of soil *R*_h_ under long-term experimental warming.

### Incorporating microbial functional gene information into ecosystem models

Due to the importance of microbes in controlling soil *R*_h_, as an exploratory effort, we further attempted to incorporate omics data into ecosystem models. Since traditional ecosystem models do not explicitly represent most microbial processes ^27^, the MEND model was employed, which explicitly represents microbial physiology and SOM decomposition catalyzed by oxidative or hydrolytic enzymes ^26^. Because MEND model requires absolute quantitative information on hydrolytic and oxidative enzymes for SOM decomposition ^26,28^, GeoChip hybridization-based data were used, which is more effective to catch the community dynamic changes as illustrated above.

The MEND model was calibrated with or without functional gene information. We referred the former to as gene amended MEND (gMEND) and the latter as traditional MEND (tMEND). We constrained gMEND by achieving the highest correlation between MEND-modeled mean annual enzyme concentrations and GeoChip-detected annual oxidative and hydrolytic gene abundances in addition to a best fit between observed and simulated *R*_h_. Our results showed high correlations (*r* = 0.74 and 0.81 for oxidative and hydrolytic enzymes, respectively) between simulated enzyme concentrations and GeoChip-detected gene abundances (Fig. S10a-b) in the control plots. Also, relatively low Mean Absolute Relative Errors (MARE = 14% and 22%, Fig. S10c-d) were also achieved between simulated and expected enzyme concentrations under warming conditions, which were the product of simulated enzyme concentrations under control and the warming-to-control ratio of GeoChip-detected gene abundances. The above modeling results indicated good agreements on the 7-year interannual variabilities between simulated enzyme concentrations and GeoChip-detected gene abundances. Furthermore, almost all of 11 model parameters were better constrained by gMEND than by tMEND (Fig. 3a and Fig. S11). The average coefficient of variation (CV) of model parameters was significantly reduced from 77% (tMEND) to 22% (gMEND) under control and from 39% (tMEND) to 17% (gMEND) under warming. In addition, the MEND-simulated *R*_*h*_ agreed well with the observed *R*_*h*_ under warming and control (Fig. 3b: R^2^ = 0.53 and 0.63, respectively). Compared to non-microbial terrestrial ecosystem model (TECO) ^29^, the MEND model improved CO_2_ efflux fitting by 5% under control and by 19% under warming (Fig. S12). Finally, the MEND-derived intrinsic *Q*_10_ values were confined from 1.20–2.42 (tMEND) to a more reasonable range of 1.27–2.13 (gMEND), as *Q*_10_ values below 2 are preferred for better global C cycle modeling ^30^. The intrinsic *Q*_10_ values also concurred with previous site-level and global-scale studies ^30,31^. The *Q*_10_ in the MEND model solely reflects the microbial responses to temperature change, which can remove confounding effects of other environmental factors. Compared to the apparent *Q*_10_ estimated by the relationship between *R*_*h*_ and soil temperature, the MEND-derived *Q*_10_ better represents the intrinsic temperature effects on microbially-mediated SOM decomposition processes, which provides a significant advance in our understanding of microbial responses to changes in temperature. Therefore, the MEND-derived intrinsic *Q*_10_ was further used to explore how much C loss is reduced by the soil microbial acclimation (*Q*_10_) under warming. Our results showed that the microbial acclimation in the warming plots would reduce 11.6±7.5% soil *R*_h_, and thus reduce soil C loss, during the 7-year experimental period, compared to the scenarios without acclimation (Fig. 4).

**Fig. 3.**
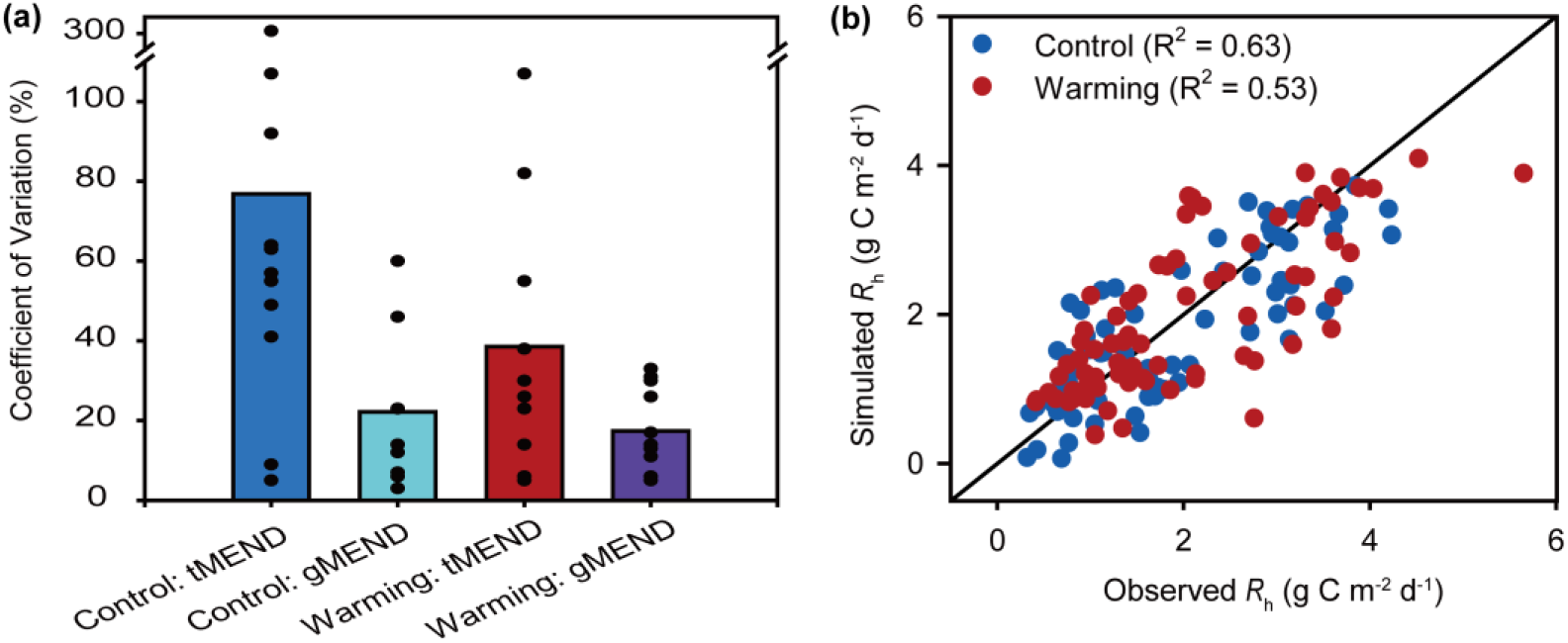
Model parameter uncertainty and modeling performance. **(a)** The MEND model parameter uncertainty quantified by the Coefficient of Variation (CV). The bars show the mean CV values of the 11 parameters (See Supplementary Fig. S11 and Table S8 for detailed description). The dots along each bar show the CV for each parameter. “tMEND” refers to the traditional MEND model parameterization without gene abundances data. “gMEND” denotes the improved MEND parameterization with gene abundances. **(b)** Comparison between gMEND-simulated and observed heterotrophic respiration (*R*_*h*_) under control and warming (R^2^ denotes the coefficient of determination).

**Fig 4.**
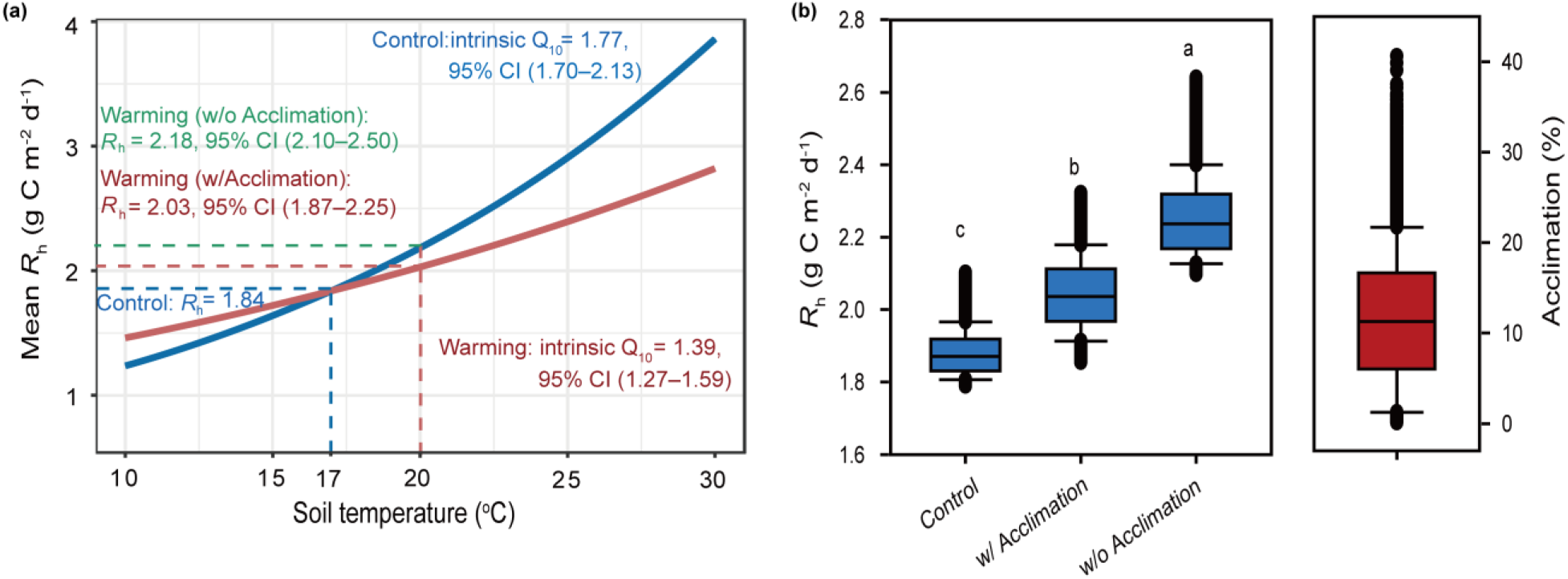
Warming-induced microbial acclimation of heterotrophic respiration (*R*_*h*_) based on MEND-estimated intrinsic *Q*_10_. **(a)** *R*_*h*_ acclimation based on the mean values. The mean annual soil temperature (*T*) during 2010–2016 was 17 °C and 20 °C under control and warming, respectively. The average intrinsic *Q*_10_ = 1.77 under control and 1.39 under warming. The mean baseline *R*_*h*_ = 1.84 g C m^−2^ d^−1^ under control (*T* = 17 °C). The average *R*_*h*_ = 2.03 and 2.18 g C m^−2^ d^−1^ under warming (*T* = 20 °C) when acclimation is considered (w/ Acclimation) or not considered (w/o Acclimation), which means a 8.2% reduction in *R*_*h*_ due to acclimation. 95% CI denotes the 95% confidence interval. **(b)** Acclimation in *R*_*h*_ when the uncertainties in intrinsic *Q*_10_ are considered. Different letters for *R*_*h*_ indicate significantly differences between the scenarios based on the Kruskal-Wallis test at a significance level of 0.05. The acclimation (%) is quantified by the difference in *R*_*h*_ between warming w/o Acclimation and w/ Acclimation as a percentage of the baseline *R*_*h*_ under control (see Methods Eq.8).

## Conclusions

Through field measurements and process model-based simulations, our results demonstrated that soil microbial respiratory acclimation persisted over the last 7 years, which is consistent with a recent long-term study on a forest ecosystem ^15^. This study provides explicit, robust evidence of the persistence of soil microbial respiratory acclimation to warming-induced rising temperature and reducing moisture over long periods. If this phenomenon holds over larger spatial scales across different ecosystems, soil microbial respiratory acclimation globally may have a greater mitigating impact than expected on climate warming-induced CO_2_ losses ^32^. If the results from this study are applicable to other grasslands globally ^33^, the microbial acclimation could lead to 0.49±0. 31 Pg (10^15^ g) less C loss per year (see Online Methods). Our study also reveals that warming-induced respiratory acclimation is significantly correlated with the adaptive changes in microbial community functional structure, which could dampen the potential positive C-climate feedbacks by reducing considerable amount of warming-induced heterotrophic respiration. In addition, although incorporating complex microbial information into global change models is extremely challenging ^20^, by parameterizing the microbial model with omics-based functional gene information, the uncertainty of key model parameters in MEND was substantially decreased, and its performance was considerably improved compared to non-microbial model. Thus, it is possible to improve the model predictive ability for projecting future environmental changes via better assessment of microbial omics-based functional capacities. However, to generalize whether these microbial mechanisms and metagenomics-enabled modeling strategy obtained in this grassland ecosystem are applicable to other ecosystems requires further long-term studies under realistic field settings.

## Materials and methods

### Site Description and Sampling

This experimental site was established in July 2009 at the Kessler Atmospheric and Ecological Field Station (KAEFS) in the US Great Plains in McClain County, Oklahoma (34° 59’ N, 97° 31’W) ^14,34^. Experimental design and site description were described in detail previously^23^. Briefly, *Ambrosia trifida*, *Solanum carolinense* and *Euphorbia dentate* belonging to C_3_ forbs, and *Tridens flavus*, *Sporobolus compositus* and *Sorghum halapense* belonging to C_4_ grasses are dominant in the site ^23,34^. Annual mean temperature is 16.3 °C and annual precipitation is 914 mm, based on Oklahoma Climatological Survey data from 1948 to 1999. The soil type of this site is Port–Pulaski–Keokuk complex with 51% of sand, 35% of silt and 13% of clay, which is a well-drained soil that is formed in loamy sediment on flood plains. The soil has a high available water holding capacity (37%), neutral pH and 1.2 g cm-3 bulk density with 1.9% total organic matter and 0.1% total nitrogen (N) ^23,34^. Four blocks were used in the field site experiment, in which warming is a primary factor. Two levels of warming (ambient and +3 °C) were set for four pairs of 2.5 m × 1.75 m plots by utilizing a “real” or “dummy” infrared radiator (Kalglo Electronics, Bethlehem, PA, USA). In the warmed plots, a real infrared radiator was suspended 1.5 m above the ground, and the dummy infrared radiator was suspended to simulate a shading effect of the device in the control plots.

In this study, eight surface (0-15 cm) soil samples, four from the warmed and four from the control plots, were collected annually at approximately the date of peak plant biomass (September or October) from 2010 to 2016. Three soil cores (2.5 cm diameter × 15 cm depth) were taken by using a soil sampler tube in each plot and composited to have enough samples for soil chemistry, microbiology and molecular biology analyses. A total of 56 soil samples were analyzed in this study.

### Environmental and soil chemical measurements

Precipitation data were obtained from the Oklahoma Mesonet Station (Washington Station) ^34^ located 200 m away from our experiment site, and 12-month version of the standardized precipitation-evapotranspiration index (SPEI-12) was used as annual drought index ^35,36^. Air temperature, soil temperature and volumetric soil water content were measured as previously described ^23^.

All soil samples were analyzed to determine soil total organic carbon (TOC), total nitrogen (TN), soil nitrate (NO_3_^−^) and ammonia (NH_4_^+^) by the Soil, Water, and Forage Analytical Laboratory at Oklahoma State University (Stillwater, OK, USA). Soil pH was measured using a pH meter with a calibrated combined glass electrode ^37^.

### Aboveground plant communities

Aboveground plant community investigations were annually conducted at peak biomass (usually September) as described previously ^34,38^. Aboveground plant biomass, separated into C_3_ and C_4_ species, was indirectly estimated by a modified pin-touch method ^34,38^. Detailed description of biomass estimation is provided by Sherry *et al*. ^39^. All of the species in plant community within each plot were identified to estimate species richness.

### Ecosystem C fluxes and soil respirations

Ecosystem C fluxes and soil respirations were measured once or twice a month between 10:00 and 15:00 (local time) from January 2010 to December 2016 as described previously 14,34. One square aluminum frame (0.5 m × 0.5 m) was inserted in the soil at 2 cm depth in each plot to provide a flat base between the soil surface and the CO_2_ sampling chamber. Net ecosystem exchange (NEE) and ecosystem respiration (ER) were measured using LI-6400 portable photosynthesis system (LI-COR) ^40^. Gross primary productivity (GPP) was estimated as the difference between NEE and ER. Meanwhile, soil surface respiration was monthly measured using a LI-8100A soil flux system attached to a soil CO_2_ flux chamber (LI-COR). Measurements were taken above a PVC collar (80 cm^2^ in area and 5 cm in depth) and a PVC tube (80 cm^2^ in area and 70 cm in depth) in each plot. The PVC tube cut off old plant roots and prevented new roots from growing inside the tube. Any aboveground parts of living plants were removed from the PVC tubes and collars before each measurement. The CO_2_ efflux measured above the PVC tubes represented heterotrophic respiration (*R*_h_) from soil microbes, while that measured above the PVC collars represented soil total respiration (*R*_t_) including heterotrophic and autotrophic respiration (*R*_h_ and *R*_a_) from soil microbes and plant root respectively.

### Soil decomposition rate

Weighted cellulose filter paper (Whatman CAT No. 1442-090) was placed into fiberglass mesh bags and placed vertically at 0-10 cm soil depth in each plot in March 2016. All of decomposition bags were collected back in September 2016, rinsed and dried at 60 °C for weighing. Th e percentage of mass loss was calculated to represent soil decomposition rate.

### Molecular analyses of soil samples

#### a. BIOLOG analysis

The C substrate utilization patterns of soil microbial communities in 2016 were analyzed by BIOLOG EcoPlate™ (BIOLOG). The BIOLOG EcoPlate™ contains 31 of the most useful labile carbon sources for soil community analysis, which are repeated 3 times in each plate. In this study, the plates with diluted soil supernatant (0.5g soil with 45 mL 0.85% NaCl) were incubated in a BIOLOG OmniLog PM System at 25 °C for 4.5 days. The color change of each well was shown as absorbance curve. The net area under the absorbance versus time curve was calculated to represent physiological activity of various C sources ^41^. The average value from 3 replicates was used for analyses in this study.

#### b. DNA extraction, amplicon sequencing and analysis

Methods for DNA extraction from soil and amplicon sequencing of all soil samples were as previously described ^23^. Briefly, 10 ng DNA per sample were used for library construction and amplicon sequencing ^42^. The V4 region of bacterial and archaeal 16S rRNA genes and fungal ITSs between 5.8S and 28S rRNA genes were amplified with primer sets 515F/806R and ITS7F/ITS4R, and sequenced on a MiSeq platform (Illumina, Inc.) using 2 × 250 pair-end sequencing kit. Raw sequences were submitted to our Galaxy sequence analysis pipeline (http://zhoulab5.rccc.ou.edu:8080) to further analyze as previously described ^23^. Finally, OTUs were clustered by UPARSE ^43^ at 97% identity for both 16S rRNA gene and ITS. All sequences were randomly resampled to 30,000 sequences for 16S rRNA gene and 10,000 sequences for ITS per sample. Representative sequences of OTUs were annotated taxonomically by the Ribosomal Database Project (RDP) Classifier with 50% confidence estimates ^44^.

#### c. GeoChip analysis

GeoChip 5.0M, a functional gene array ^45^, was used for all 56 samples from 2010 to 2016. GeoChip hybridization, scanning and data processing were performed as described previously ^45^. The raw signals from NimbleGen were submitted to the Microarray Data Manager on our website (http://ieg.ou.edu/microarray), cleaned, normalized and analyzed using the data analysis pipeline. Briefly, spot signal-to-noise ratio and minimum intensity cutoff were used as standard to remove unreliable spots. Both the universal standard and functional gene spot intensities were used to normalize the signals among arrays ^45^.

#### d. Shotgun sequencing analysis

Metagenomic library of all samples was prepared using a KAPA Hyper Prep Kit and sequenced at the Oklahoma Medical Research Foundation’s Genomics Core using the Illumina HiSeq 3000 platform with a 2 × 150 bp paired-end kit. A total of 8.18 billion reads were obtained from all 56 samples, and 80 million reads were randomly resampled from each sample to perform data processing. Functional gene prediction, annotation and treatment analyses were performed using methods similar to those described in previous study ^45^. Meanwhile, all reads were also submitted to our EcoFUN-MAP pipeline (http://www.ou.edu/ieg/tools/data-analysis-pipeline.html) to fish out shotgun sequence reads of important environmental functional genes used to fabricate GeoChip as described previously ^46^. The web based pipeline application of EcoFUN-MAP can be accessed with request.

### Model simulations (TECO and MEND model)

#### a. Data sources

Daily GPP values were obtained from a corrected 8-day GPP product based on the MODIS GPP (MOD17A2/MOD17A2H) ^47^. We assign the same daily GPP values for the 8-day period. Meanwhile, data sets measured in both control and warmed plots across all years were also used for model simulations, including soil temperature and moisture, heterotrophic respiration, and the GeoChip-detected enzyme densities.

#### b. Apparent Q_10_ estimation

To examine temperature sensitivity of microbial heterotrophic respirations, the measured field *R*_h_ in warmed and control plots was fitted with the exponential equation ^4^ (Equation (1)) on yearly basis or across all years. In the equation, *R* is *R*_h_, *T* is soil temperature, *R*(*T*_*ref*_) is the respiration rate at the reference temperature (*T*_*ref*_). The *Q*_10_ estimated by the observed respiration data was called apparent *Q*_10_ of respiration in this study.

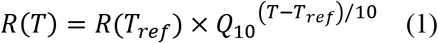

#### c. Intrinsic *Q*_10_ estimation

In the MEND model, the parameter *Q*_10_ is used to characterize the unconfounded temperature sensitivity of SOM decomposition and heterotrophic respiration. Constrained *Q*_10_ were obtained for the control and warming plots by incorporating respiration and microbial information into the MEND model parameterization process, which we called the intrinsic *Q*_10_ of soil respirations ^30^.

#### d. TECO model

The non-microbial terrestrial ecosystem (TECO) model is a variant of the CENTURY model ^48^ that is designed to simulate C input from photosynthesis, C transfer among plant and soil pools, and respiratory C releases to the atmosphere (Fig. S3b). C dynamics in the TECO model can be described by a group of first-order ordinary differential equations, where the turnover rates are modified by soil temperature (*T*) and moisture (*W*) ^29^. Prior ranges of turnover rates were based on Weng and Lu ^49^. The prior ranges of *Q*_10_ were based on the ranges of apparent *Q*_10_ of *R*_h_ per treatment ^4^. We assumed that the parameters distributed uniformly in their *prior* ranges ^8^. We used the Shuffled Complex Evolution (SCE) algorithm to determine model parameters ^26^. We also applied the probabilistic inversion (Markov Chain Monte Carlo) to quantity parameter uncertainties ^50^. By performing TECO modeling, daily heterotrophic respiration was simulated for both warmed and control plots from 2010 to 2016. The coefficient of determination (*R*^2^) was used to estimate the model performance between observed and simulated respiration ^51^.

#### e. Microbial-ENzyme Decomposition (MEND) model

##### e.1. MEND model description

The Microbial-ENzyme Decomposition (MEND) model (Fig. S3a) describes the SOM decomposition processes by explicitly representing relevant microbial and enzymatic physiology ^26^. The SOM pool consists of two particulate organic matter (POM) pools and one mineral-associated organic matter (MOM) pool. The two POMs are decomposed by oxidative and hydrolytic enzymes, respectively. The MOM is decomposed by a generic enzyme group (EM). Model state variables, governing equations, component fluxes and parameters are described in Table S6–S9, respectively. A model parameter (reaction rate) in MEND may be modified by soil water potential, temperature, or pH ^26,52^. MEND represents microbial dormancy, resuscitation, and mortality and enzymatic decomposition in response to changes in moisture, as well as shifting of microbial and enzymatic activities with changing temperature ^25^. The temperature response functions are described by the Arrhenius equation (characterized by the activation energy) or the *Q*_10_ method ^53^, where the *Q*_10_ method was used in this study.

##### e.2. Model Parameterization

The model parameters are determined by achieving high goodness-of-fits of model simulations against experimental observations, such as heterotrophic respiration (*R*_h_), microbial biomass carbon (MBC), gene abundances of oxidative (EnzCo) and hydrolytic enzymes (EnzCh) in this study (Table S10). We implemented multi-objective calibration of the model ^25^. Each objective evaluates the goodness-of-fit of a specific observed variable, e.g., *R*_h_, MBC, or gene abundances (Table S10). Note that the GeoChip gene abundances were used to constrain the MEND modeling as additional objective functions. The parameter optimization is to minimize the overall objective function (*J*) that is computed as the weighted average of multiple single-objectives (Table S9) ^26^

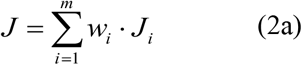

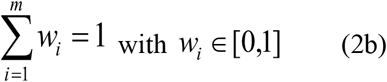

Where *m* denotes the number of objectives and *w*_*i*_ is the weighting factor for the *i*^th^ (*i* = 1,2,…,*m*) objective (*J*_*i*_). In this study, *J*_*i*_ (*i*=1, 2, 3, 4) refers to the objective function value for *R*_h_, MBC, EnzCo, and EnzCh, respectively.

As the overall objective function *J* is minimized in the parameter optimization process, the individual objective function *J*_*i*_ may be calculated as (1− *R*^2^), (1−*r*), or *MARE*:

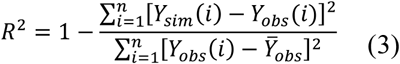

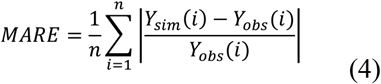

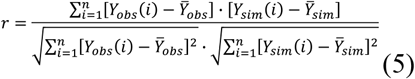

where *R*^2^ denotes the Coefficient of Determination ^26,54^. The R^2^ quantifies the proportion of the variance in the response variables that is predictable from the independent variables. A higher R^2^ (*R*^2^ ≤ 1) indicates better model performance. *MARE* is the Mean Absolute Relative Error (MARE) and lower *MARE* values (*MARE* ≥ 0) are preferred ^26,55^. *MAR*E represents the averaged deviations of predictions (*Y*_*sim*_) from their observations (*Y*_*obs*_). *r* is Pearson correlation coefficient and higher *r* values (|*r*|≤ 1) means better model performance. *n* is the number of data; *Y*_*obs*_ and *Y*_*sim*_ are observed and simulated values, respectively; and 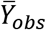 and 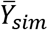 are the mean value for *Y*_*obs*_ and *Y*_*sim*_, respectively.

Different objective functions are used to quantify the goodness-of-fit for different variables (Table S9), depending on the measurement method and frequency of variables. The R^2^ is used to evaluate the variables (e.g., soil respiration) that are frequently measured and the absolute values can be directly compared between observations and simulations. The MARE is used to evaluate the variables (e.g., microbial biomass and enzyme concentrations) with only a few measurements and the absolute values can be directly compared. When the absolute values cannot be directly compared, the correlation coefficient (*r*) between original or transformed (e.g., logarithmic transformed) observations and simulations will be used. For example, the gene abundances from metagenomics or GeoChip analysis cannot be directly compared to the enzyme concentrations or activities in the MEND model. However, we may assume correlation could be found between the measured and modeled values with a certain transformation or normalization.

We used the Shuffled Complex Evolution (SCE) algorithm to determine model parameters for the control soil and the warming soil respectively. SCE is a stochastic optimization method that includes competitive evolution of a ‘complex’ of points spanning the parameter space and the shuffling of complexes ^56^.

##### e.3. Uncertainty quantification

The parameter uncertainty in the MEND model was quantified by the Critical Objective Function Index (COFI) method ^26,52^. The COFI method is based on a global stochastic optimization technique (e.g., SCE in this study). It also accounts for model complexity (represented by the number of model parameters) and observational data availability (represented by the number of observations). The confidence region of parametric space were determined by selecting those parameter sets resulting in objective function values (*J*) less than the COFI value (*J*_*cr*_) from the feasible parameter space ^26,52^.

##### e.4. Estimation of warming-induced soil C loss and acclimation effect

To examine how much soil C loss is reduced by the soil microbial respiratory acclimation under warming, we further calculated heterotrophic respiration (*R*_*h*_) under warming without acclimation (w/o Acclimation). That is, we estimated the mean *R*_*h*_ changing with soil temperature that under warming, however, we kept the same range of *Q*_10_ as that under control ^13,15^. The *R*_*h*_ changing with soil temperature is described by the *Q*_10_ method similar to Eq. (1):

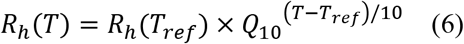

where *R*_*h*_(*T*) and *R*_*h*_(*T*_*ref*_) are the *R*_*h*_ (g C m^−2^ d^−1^) at soil temperature (*T*) and reference temperature (*T*_*ref*_), respectively; and *T*_*ref*_ = 10 °C in this study.

We quantified the acclimation effect by taking into account the uncertainties in intrinsic *Q*_10_ estimated by the MEND model. First we calculated the *R*_h_ fluxes (g C m^−2^ d^−1^) at the mean annual soil temperature under control, i.e., 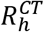 under *T* = 17 °C and *Q*_10_ = 1.77 with 95% confidence interval (CI) of 1.70–2.13. Second we calculated *R*_*h*_ under warming with acclimation (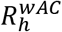 under *T* = 20 °C and *Q*_10_ = 1.39 with 95% CI of 1.27–1.59) and *R*_*h*_ under warming without acclimation (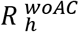 under *T* = 20 °C and *Q*_10_ = 1.77 with 95% CI of 1.70–2.13). We then calculated the reduction in *R*_*h*_ due to acclimation as

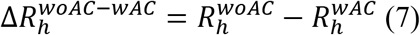

Finally, we calculated the acclimation effect as the percent reduction in *R*_*h*_ due to acclimation relative to the baseline *R*_*h*_, i.e, the mean *R*_*h*_ in the control plot 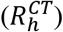

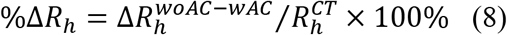

As a preliminary test of global significance, we extrapolated our results to the world’s grasslands: The annual soil respiration flux (*R*_*s*_) was 8.0 Pg C yr^−1^ in the global grasslands (area = 1.11×10^7^ km^2^) with the MODIS land cover map in 2009 according to Adachi et al. ^33^, which meant *R*_*s*_ = 720.7 g C m^−2^ yr^−1^.

We then estimated the heterotrophic respiration flux *R*_*h*_ = 381.7 gC m^−2^ yr^−1^ and *R*_*h*_ / *R*_*s*_ = 53% in global grasslands based on the relationship between *R*_*h*_ and *R*_*s*_ (in units of g C m^−2^ yr^−1^) described by Bond-Lamberty & Thomson ^57^:

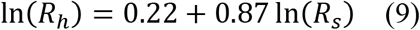

Based on the ratio *R*_*h*_ / *R*_*s*_ = 53%, the annual heterotrophic respiration flux from global grasslands was estimated as 4.2 Pg C yr^−1^.

We then estimated the acclimation effect as follows. Our results show that %Δ*R*_*h*_= 11.6±7.5% (Fig. 4c), which means the reduction in *R*_*h*_ due to acclimation accounted for 11.6±7.5% of 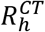. If this percentage is applicable to the global grasslands, the warming acclimation would result in less *R*_*h*_ by 0.49±0.31 Pg C yr^−1^ (= 4.2 Pg C yr^−1^×(11.6±7.5%)).

### Statistical analysis

All statistical analyses were carried out using R software 3.1.1 with the package vegan ^58^ (v.2.3-5) and pgirmess ^59^ (v.1.5.8) unless otherwise indicated. The difference of various variables between warming and control was tested by repeated-measures analysis of variance (ANOVA). The non-parametric multivariate analysis of variance (Adonis) were used to test the difference of microbial community taxonomic and functional structures considering the blocked split-plot design ^23^. CCA and Mantel test were performed to examine the linkage between environmental variables and microbial community structure/subcategories of functional genes. The significance of the CCA model was tested by analysis of variance (ANOVA). CCA-based variation partitioning analysis (VPA) was performed to evaluate how much different types of environmental variables influences microbial community phylogenetic and functional structures ^14^. Structural equation model (SEM) was used to explore how warming-induced environmental variables affected soil microbial communities and heterotrophic respiration. Response ratio was used to compute the effects of warming on functional genes involved in C cycling and nutrient-cycling processes from GeoChip data using the formula as previously described ^46^. The non-parametric Kruskal-Wallis method ^59^ was used to test the significance of difference in model parameter values or the *R*_*h*_ under different scenarios at a significance level of 0.05.

## Data and code availability

DNA sequences of 16S rRNA gene and ITS amplicons were available in NCBI Sequence Read Archive under project no. PRJNA331185. Raw shotgun metagenomic sequences are deposited in the European Nucleotide Archive (http://www.ebi.ac.uk/ena) under study no. PRJNA533082. GeoChip signal intensity data can be accessed through the URL (https://www.ou.edu/ieg/publications/datasets). MEND model code and data are accessible at https://github.com/wanggangsheng/MENDokw.git. All other relevant data are available in Supplementary Information or from the corresponding author upon request.

## Acknowledgement

We are grateful to the numerous former laboratory members for their help in maintaining the experimental site. We thank Drs. Xiangming Xiao, Xiaocui Wu, and Rajen Bajgain at University of Oklahoma for providing data support. This study is funded by the US Department of Energy, Office of Science, Genomic Science Program under Award Number DE-SC0004601 and DE-SC0010715, and the Office of the Vice President for Research at the University of Oklahoma. X.G., Q.G. and X.Z. acknowledge China Scholarship Council (CSC) for support.

## Author contributions

All authors contributed intellectual input and assistance to this study and manuscript preparation. Research questions and experimental strategy were developed by J.Z., E.A.G.S., Y.L., K.T.K., J.R.C., C.R.P., and J.M.T. Field management was carried out by J.F., M.Y., C.G.J., S.N., D.L., X.X., L.J., LY.W., A.Z., F.L., B.W. and J.D.V.N. Sampling collections, DNA preparation and MiSeq sequencing analysis were carried out by X.G., L.C., and X.Z. GeoChip hybridization and shotgun sequencing analysis were performed by X.G., X.Z., and R.T. Soil chemical and substrate analyses were carried out by X.Z., L.H., A.E. and LW.W. Modeling was done by Q.G., and G.W. Various statistical analyses were carried by X.G., Z.S., and D.N. Assistance in data interpretation was provided by X.L., Y.Y., and Z.H. All data analysis and integration were guided by J.Z. The paper was written by J.Z., X.G., and Q.G. with help from G.W. Considering their contributions in terms of site management, data collection, analyses and/or integration over the last 8 years, X.G., Q.G., M.Y., and G.W. are listed as co-first authors.

