## Supplementary information for "Lower Soil Carbon Loss Due to Persistent Microbial Adaptation to Climate Warming"

### Supplementary Figures and Tables

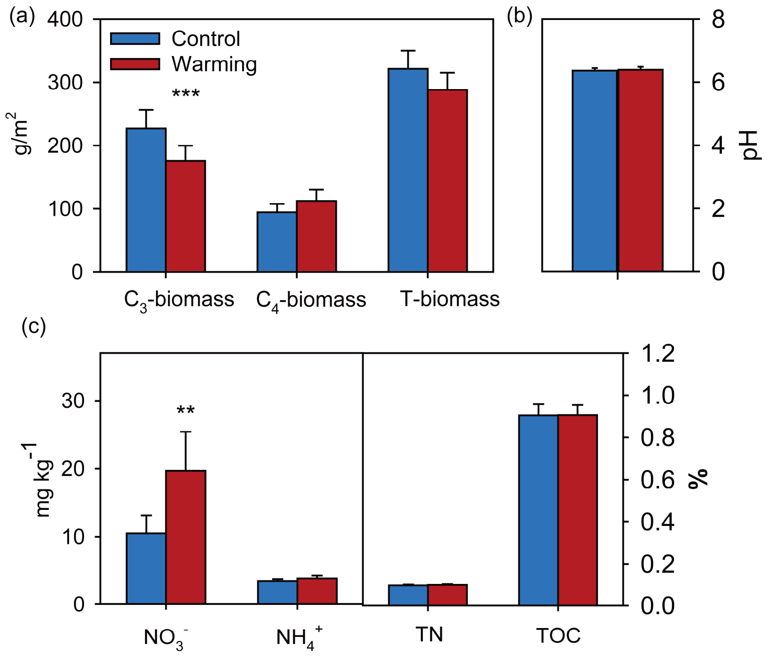


**Fig. S1. Warming effects on plant and soil variables. (a)** Effects of warming on aboveground plant biomass from $C_3$, $C_4$ and total species; (**b**) Soil pH; (**c**) Soil nitrate ($NO_3^-$), ammonia ($NH_4^+$), total N (TN) and total organic carbon (TOC) across 7 years. Error bars represent standard error of the mean ($n$ = 28). The differences between warming and the control were tested by repeated-measures ANOVA, indicated by *** when p < 0.01, ** when p < 0.05.

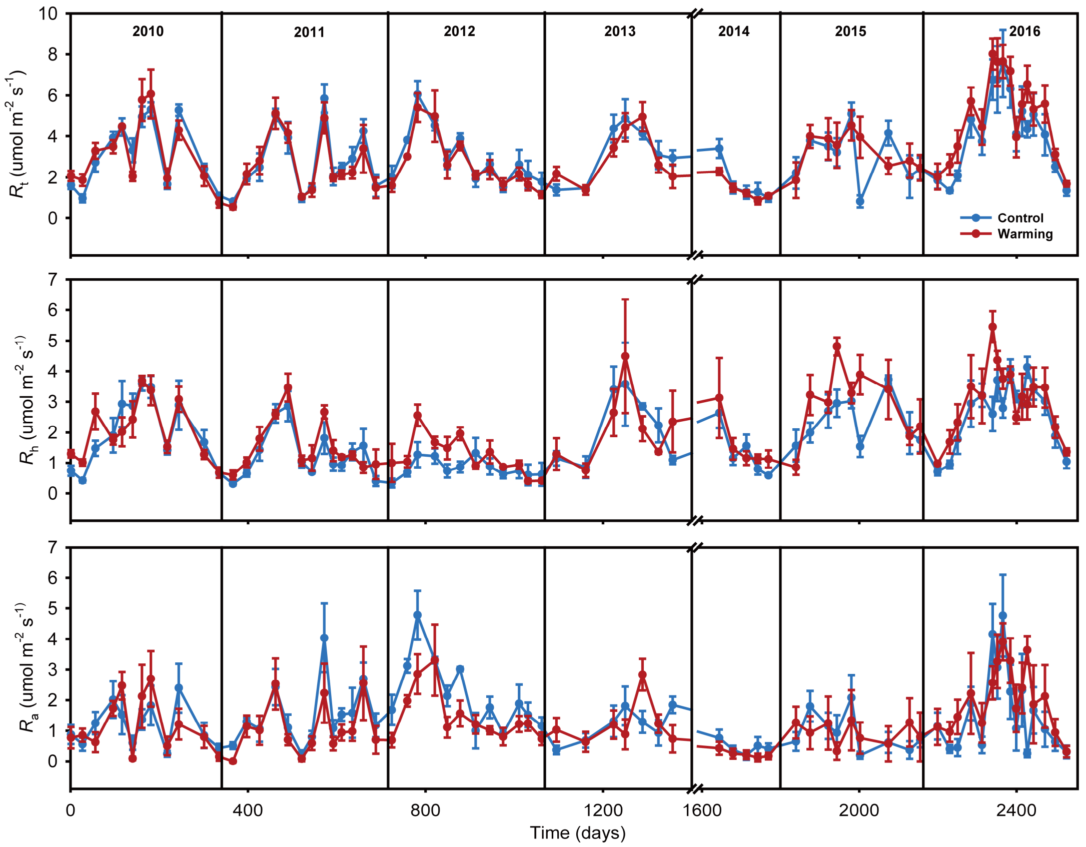


**Fig. S2. Temporal change of soil respiration ($R_t$), heterotrophic respiration ($R_h$) and autotrophic respiration ($R_a$) from 2010 to 2016**. The respiration values were displayed as mean ± standard error ($n$ = 4).

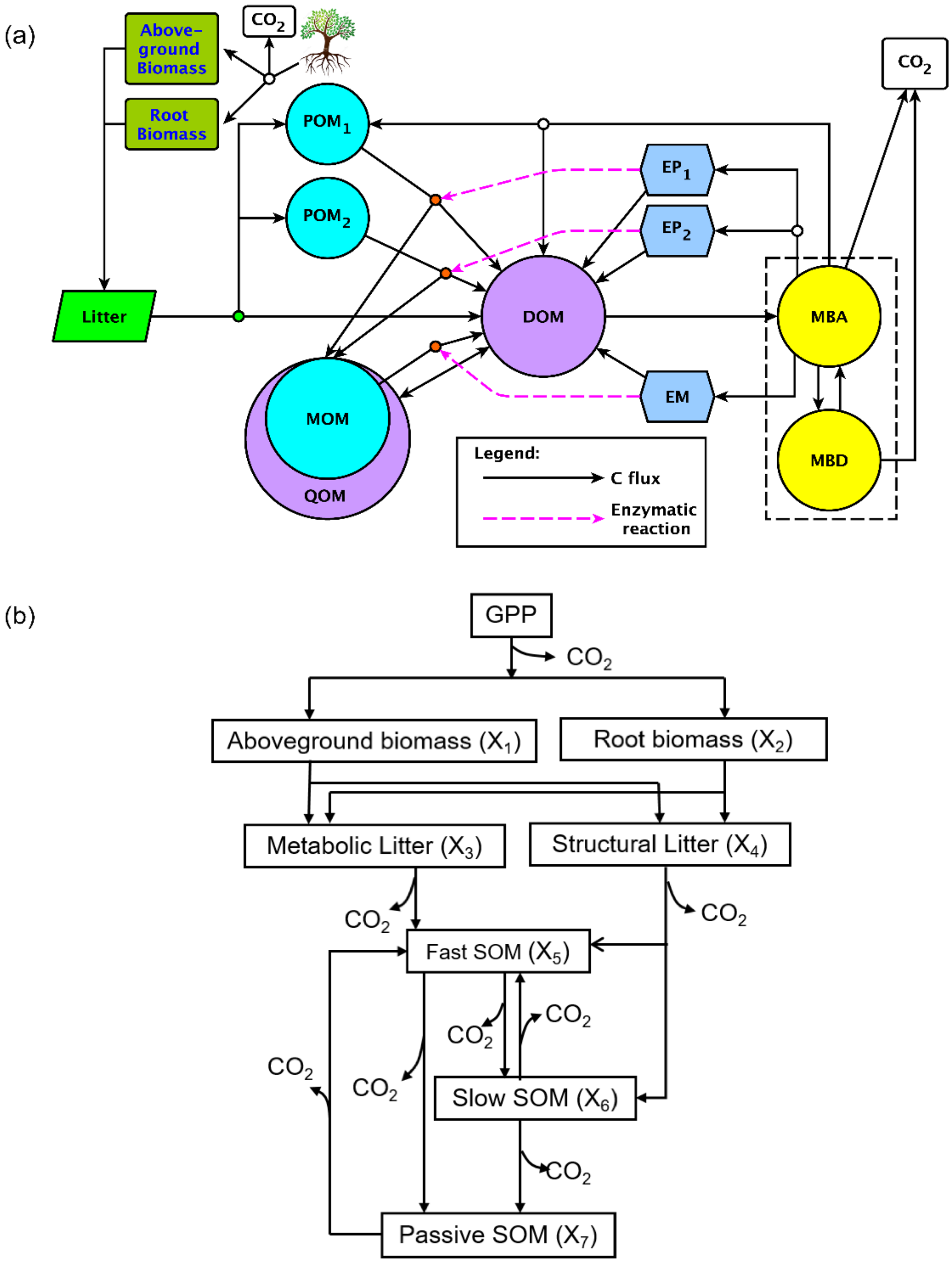


**Fig. S3. Flowcharts of ecosystem models. (a)** Microbial-ENzyme Decomposition (MEND) model. Soil organic matter (SOM) pools include: particulate organic matter (POM) (e.g., POM decomposed by oxidative and hydrolytic enzymes, denoted by $P_1$ and $P_2$ in the governing equations, respectively), mineral-associated organic matter (MOM, denoted by $M$), dissolved organic matter (DOM, $D$), adsorbed phase of DOM (QOM, $Q$), active and dormant microbes (MBA and MBD, denoted by $BA$ & $BD$), POM-degraded enzymes (e.g., $EP_1$ and $EP_2$ that break down $P_1$ and $P_2$, respectively), and MOM-degraded enzymes ($EM$). **(b)** Terrestrial ECOsystem (TECO) model.

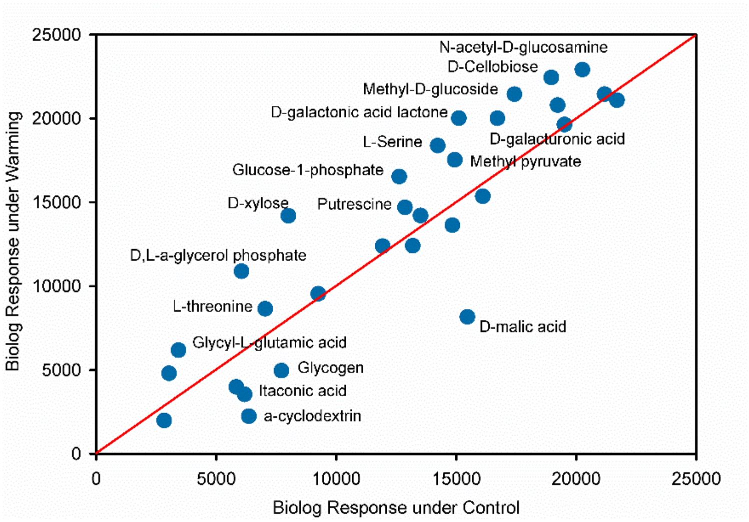


**Fig. S4. A scatterplot of BIOLOG metabolic profiles under warming and control in 2016.** Values close to the reference line (red) are in good agreement with the control values. Values above the reference line have an enhanced ability to utilize that carbon source in the warmed plots, value below have an inhibited ability in the warmed plots.

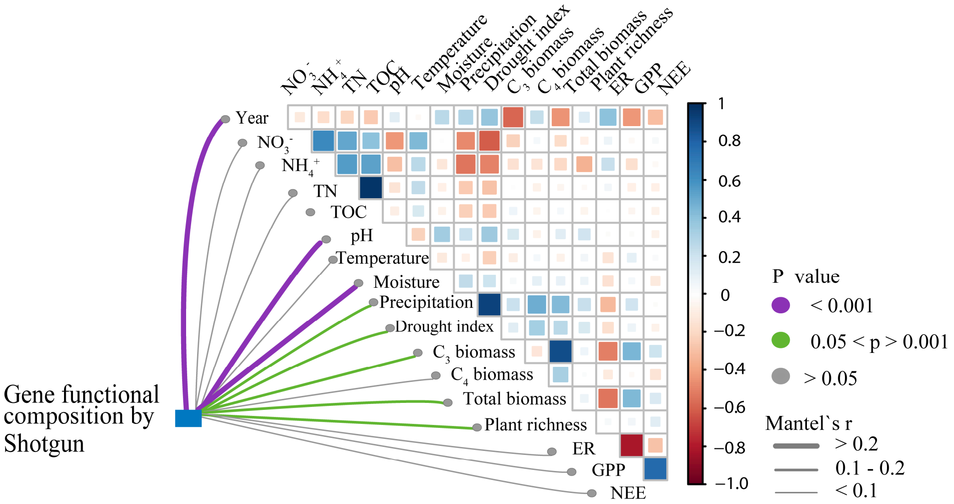


**Fig. S5. Pairwise comparisons of environmental factors with functional community structure based on shotgun sequencing data.** The shotgun sequencing data were annotated using EcoFUN-MAP database. A color gradient denotes Pearson's correlation coefficients with functional community structure by partial Mantel tests. Edge width corresponds to the Mantel's *r* statistic for the corresponding distance correlations, and edge color denotes the statistical significance.

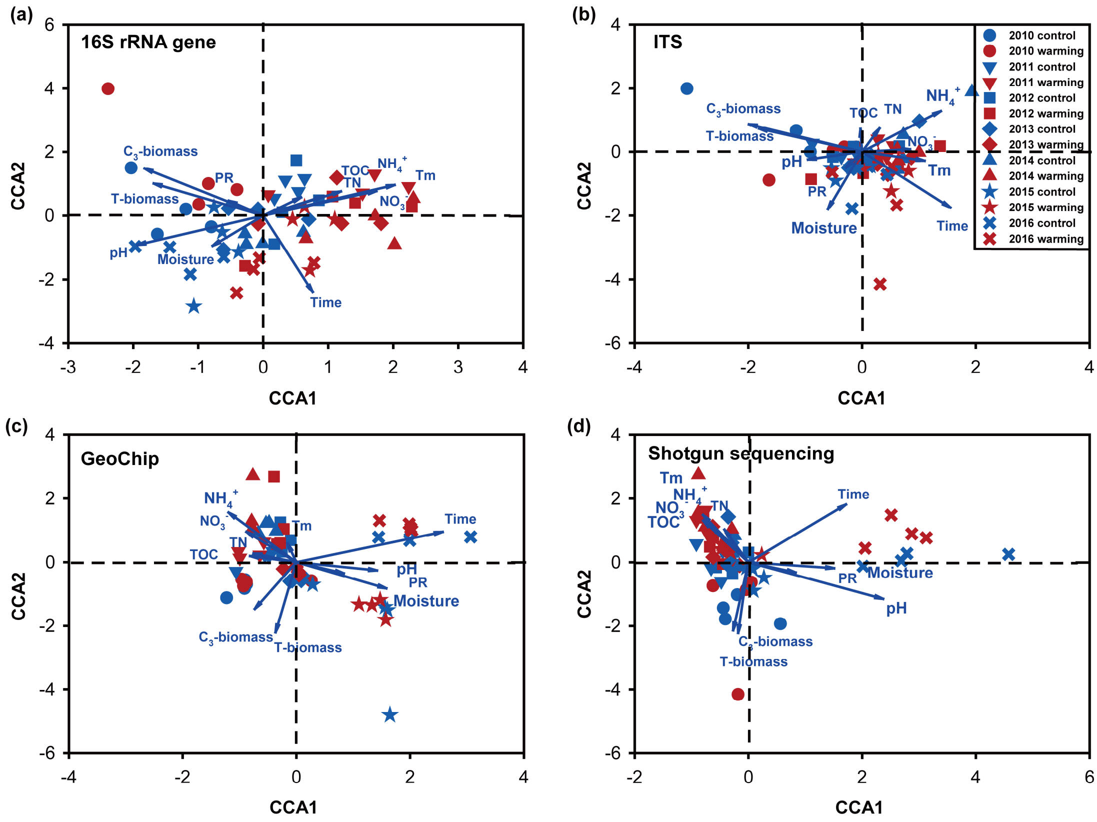


**Fig. S6. Canonical correspondence analyses (CCA) of microbial communities. (a)** Bacterial community based on 16S rRNA gene; **(b)** Fungal community based on ITS; **(c)** Functional community based on GeoChip; and **(d)** Functional community based on shotgun metagenomic sequences with EcoFUN-MAP. Phylogenetic and functional structures of microbial communities were significantly shaped by soil related factors: soil temperature (Tm), moisture, soil pH, soil total organic carbon (TOC), total nitrogen (TN), soil nitrate ($NO_3^-$) and ammonia ($NH_4^+$) contents; by plant related factors: $C_3$ and total aboveground plant biomass, and plant richness (PR); and by time.

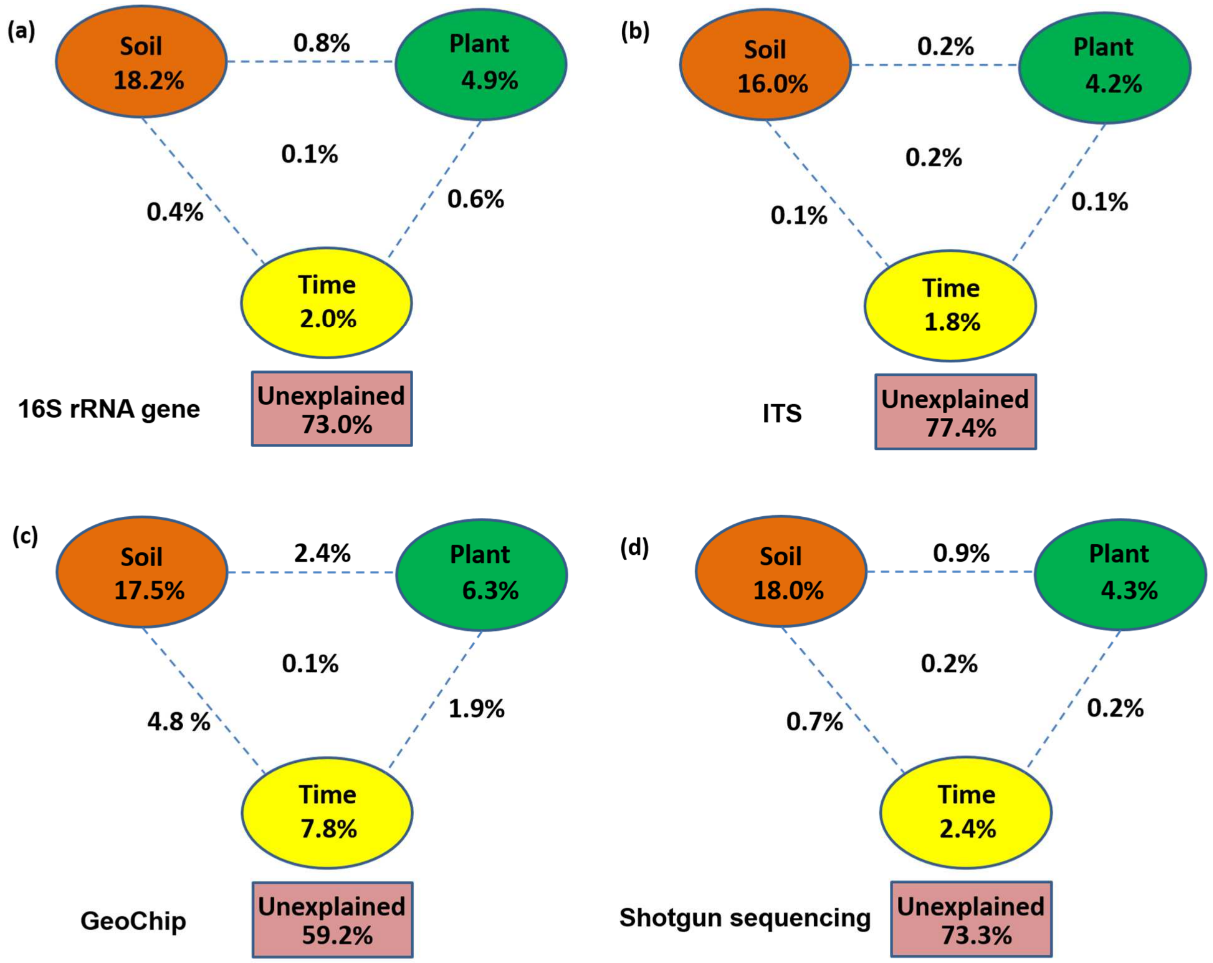


**Fig. S7. CCA-based variation partitioning analysis (VPA) of microbial communities. (a)** Bacterial community based on 16S rRNA gene; **(b)** Fungal community based on ITS; **(c)** Functional community based on GeoChip; and **(d)** Functional community based on shotgun metagenomic sequences based on EcoFUN-MAP. The relative proportions of bacterial community variations that can be explained by different types of environmental factors including soil related factors: soil temperature (Tm), moisture, soil pH, soil total organic carbon (TOC), total nitrogen (TN), soil nitrate ($NO_3^-$) and ammonia ($NH_4^+$) contents; plant related factors: $C_3$ and total aboveground plant biomass, and plant richness (PR); and time. The unexplained variations are either due to unmeasured environmental variables and/or stochastic factors.

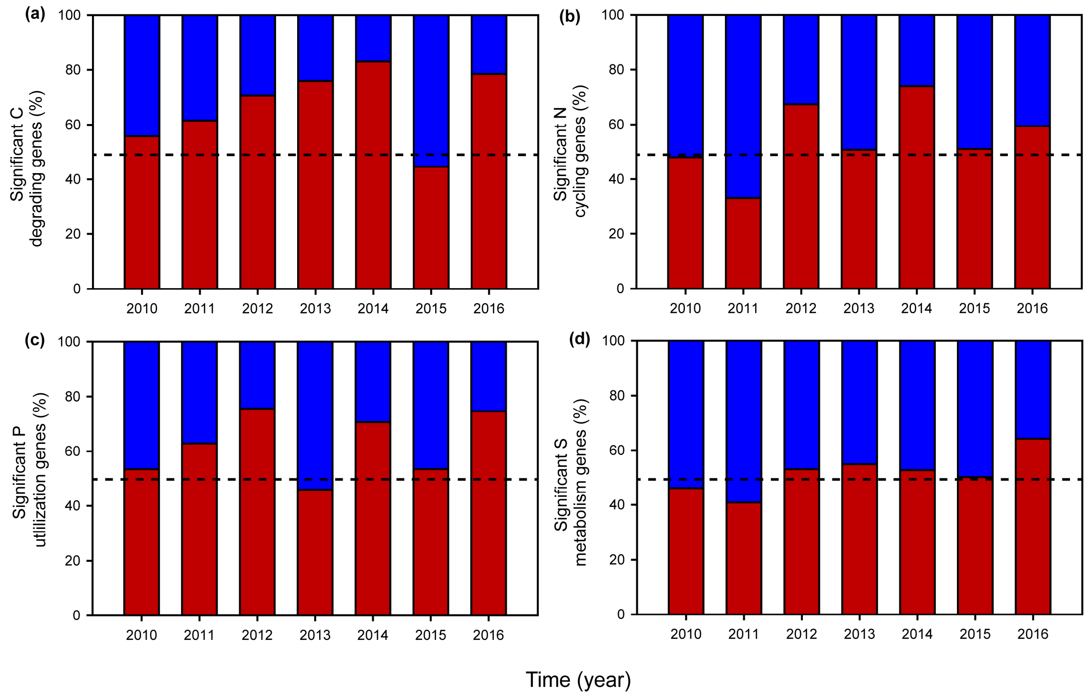


**Fig. S8. Signfiicantly changed genes involoved in C degradation (a), N cycling (b), P utilization (c) and S metabolism (d) by warming according to GeoChip data.** Significance is based on response ratio of each gene with 95% confidence intervals of abundance differences between warmed and control treatments. Dash line represents that the abundance of warming-stimulated (red) genes are in good agreement with the abundance of warming-inhibited (blue) genes. The genes involved in C degradation, N cycling, P utilization and S metabolism in this plot are listed in Supplementary Table S5

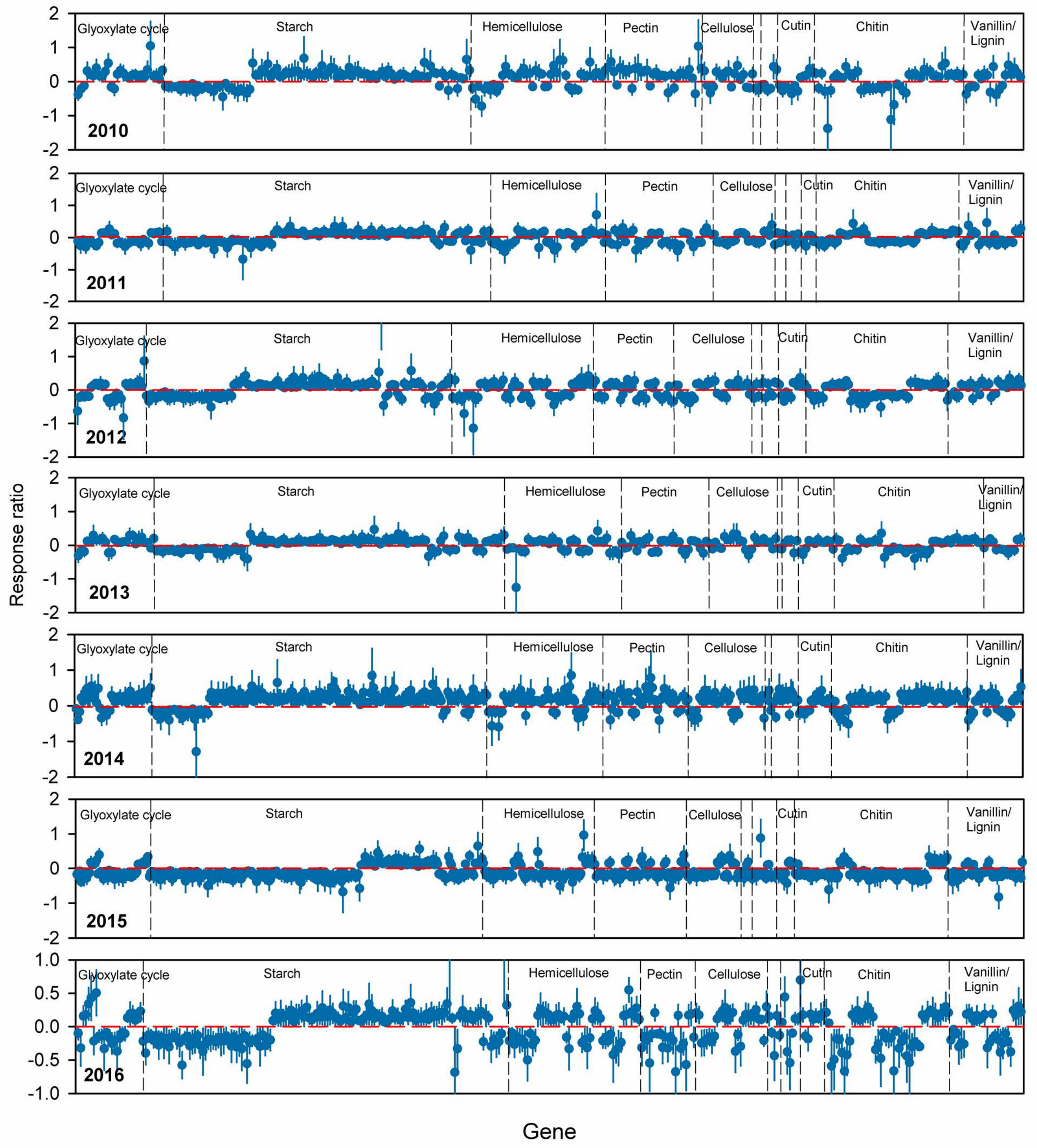


**Fig. S9. Response ratios showing significant changes in abundance of C degradation genes in each year detected by GeoChip**. Warming-stimulated C degrading genes were more than warming- inhibited genes in most years. Error bars represented 95% confidence intervals of abundance differences between warmed and control treatments. The targeted substrates were arranaged in order from labile to recalcitrant C. The full names of the genes in this figure are listed in Supplementary Table S5.

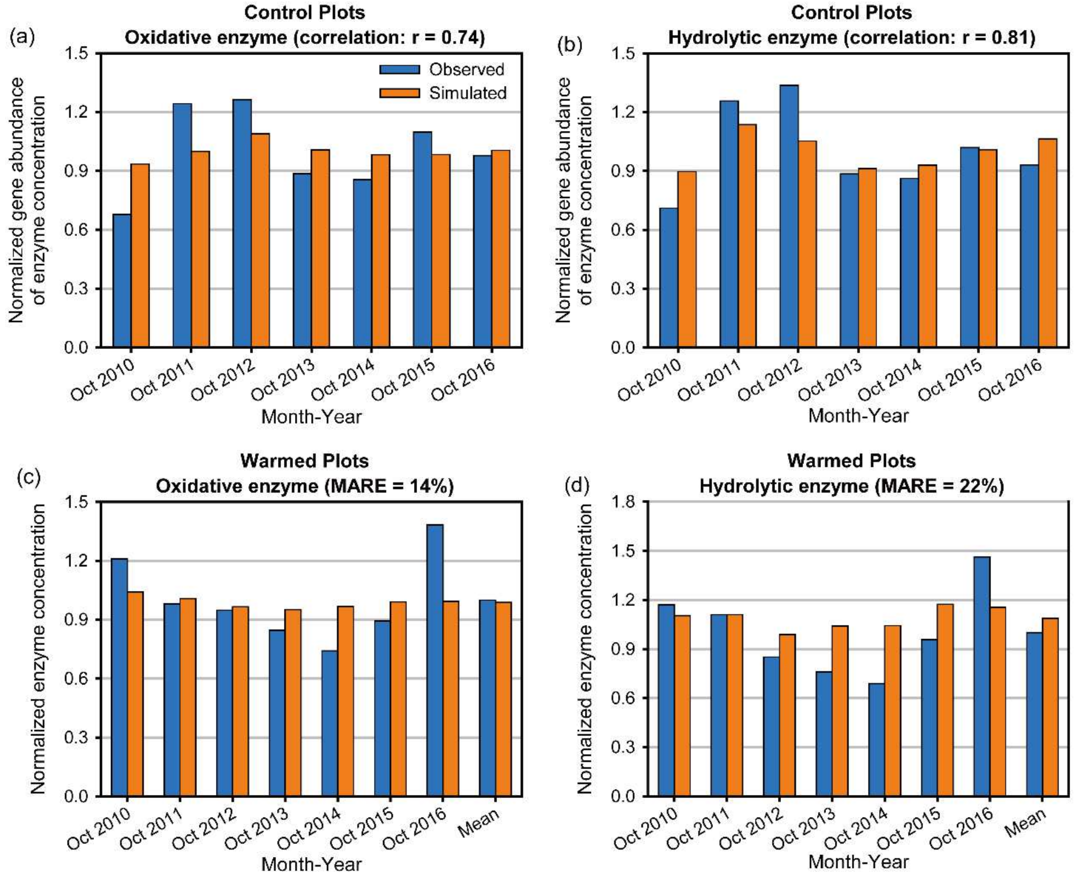


**Fig. S10. MEND modeling performance with gene abundance data.** MEND-simulated enzyme concentrations vs. GeoChip gene abundances for (**a**) oxidative enzymes and (**b**) hydrolytic enzymes in the control plot. MEND-simulated enzyme concentrations vs. GeoChip-informed enzyme concentrations for (**c**) oxidative enzymes and (**d**) hydrolytic enzymes in the warmed plot. The model performance for the control plot is quantified by the correlation coefficient (*r*), as we cannot directly compare the absolute values between GeoChip gene abundances and MEND enzyme concentrations. The model performance for the simulations under warming is evaluated by the Mean Absolute Relative Error (*MARE*, ≥0%) (see Table S9). Lower *MARE* value means better performance. All data are normalized by their respective mean values.

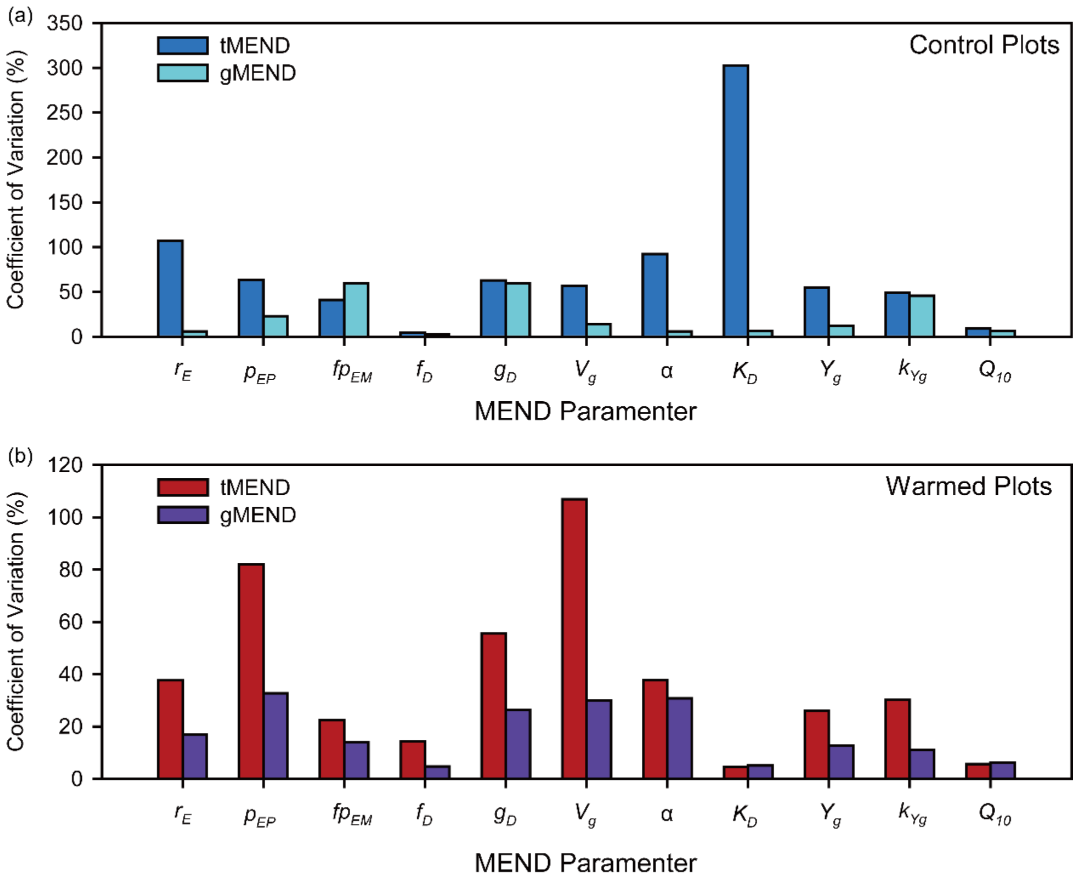


**Fig. S11. The MEND model parameter uncertainty was quantified by the Coefficient of Variation (CV) in the (a) Control and (b) Warmed plot.** "tMEND" refers to the traditional MEND model parameterization without gene abundances data. "gMEND" denotes the improved MEND parameterization with gene abundances. The 11 model parameters are $r_E$: enzyme turnover rate; $p_{EP}$ and $fp_{EM}$: two coefficients controlling enzyme production rates; $f_D$: fraction of decomposed particulate organic matter (POM) entering dissolved organic matter (DOM) pool; $g_D$: fraction of dead microbe entering DOM pool; $V_g$: maximum specific growth rate for microbe; $\alpha$: a coefficient relating specific microbial maintenance rate ($V_m$) to growth rate ($\alpha = V_m / (V_g + V_m)$); $K_D$: half-saturation constant for microbial uptake of DOM; $Y_g$: intrinsic carbon use efficiency at reference temperature; $k_{Yg}$: temperature sensitivity of $Y_g$; $Q_{10}$: temperature sensitivity of enzyme-catalyzed soil organic matter decomposition. See Table S9 for detailed description of all model parameters.

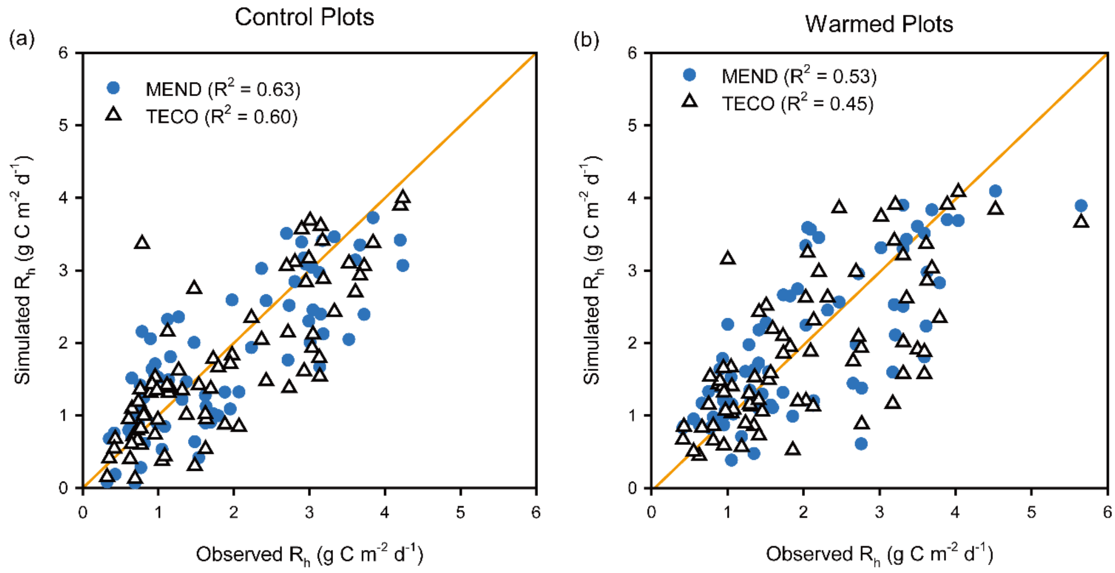


**Fig. S12. Improvement of model performance with MEND compared to the non-microbial model TECO**. (**a**) Control plots. (**b**) Warmed plots.

**Table S1.** Apparent temperature sensitivity of soil heterotrophic respiration ($Q_{10}$) by the curve fitting method for the control and warmed treatemnts in each year (2010–2016).

| | Control | | | Warming | | |
|---|---|---|---|---|---|---|
| Year | $Q_{10}$ | $R^2$ | p | $Q_{10}$ | $R^2$ | p |
| 2010 | 1.78 | 0.48 | **<0.01** | 1.58 | 0.40 | **<0.01** |
| 2011 | 1.11 | 0.02 | 0.32 | 1.12 | 0.04 | 0.14 |
| 2012 | 1.29 | 0.08 | **0.04** | 1.16 | 0.06 | **0.05** |
| 2013 | 1.60 | 0.18 | **0.02** | 1.59 | 0.10 | **0.09** |
| 2014 | 1.72 | 0.38 | **<0.01** | 1.28 | 0.25 | **0.02** |
| 2015 | 1.57 | 0.23 | **<0.01** | 1.48 | 0.22 | **<0.01** |
| 2016 | 1.66 | 0.36 | **<0.01** | 1.35 | 0.16 | **<0.01** |

**Table S2.** Summary of sequence and GeoChip statistics. The microbial samples from each year were analyzed with various molecular approaches.

| Sequencing/ GeoChip | Targets | Numbers of samples analyzed | Total base pairs (bp) | Average No. of reads/probes per sample | OTUs/ genes |
|---|---|---|---|---|---|
| 16S rRNA gene amplicon sequencing | Bacteria + archaea | 56 | 0.74G | 51,415±2,696 | 26,158 |
| ITS amplicon sequencing | Fungi | 56 | 0.43G | 3,1203±4,017 | 5,336 |
| Shotgun sequencing | Functional genes | 56 | 0.96T | 127.83±2.89M | 98,682 |
| GeoChip | Functional genes | 56 | NA | 35,425±468 | 35,425 |

**Table S3.** The correlations between the structure of each functional gene group involved soil C decomposition and N cycling processes and each environmental attribute revealed by CCA analysis. The functional community structure was determined by deep metagenome sequencing. Significance is represented by *** when p < 0.01, ** when p <0.05 and * when p <0.10.

| | | GeoChip | | | Metagenome based EcoFUN-MAP | | | Metagenomic sequencing | | |
|---|---|---|---|---|---|---|---|---|---|---|
| | Attributes+ | $R_h$ | $R_t$ | $Q_{10}$ | $R_h$ | $R_t$ | $Q_{10}$ | $R_h$ | $R_t$ | $Q_{10}$ |
| | Carbon cycling | *** | *** | | * | *** | | * | *** | ** |
| | Carbon degradation | *** | *** | * | * | ** | | ** | * | ** |
| **C degradation** | Starch | *** | *** | * | | * | | ** | * | |
| | Pectin | *** | *** | * | | | | | | ** |
| | Hemicellulose | *** | * | | | * | | | * | ** |
| | Cellulose | *** | *** | * | | | | * | * | |
| | Chitin | *** | *** | * | | * | | *** | * | |
| | Other | *** | *** | * | * | *** | | * | ** | * |
| | Vanillin/lignin | *** | *** | * | * | * | ** | | ** | ** |
| **N cycling** | Ammonification | *** | *** | * | * | ** | * | *** | *** | ** |
| | Anammox | | | | | | | *** | *** | |
| | Assimilatory N reduction | *** | *** | * | * | *** | * | | *** | ** |
| | Denitrification | *** | ** | | | | | | ** | |
| | Dissimilatory N reduction | | | | | * | | | | ** |
| | Nitrification | *** | *** | ** | | | | *** | *** | |
| | Nitrogen fixation | *** | *** | * | | | | | | |
| **P utilization** | P utilization | *** | *** | * | * | * | * | *** | *** | ** |
| **S metabolism** | Adenylylsulfate reductase | *** | *** | * | | | | | ** | * |
| | Sulfur assimilation | *** | *** | * | *** | *** | | | | |
| | Sulfite reduction | *** | *** | *** | | | | ** | *** | *** |
| | Sulfide oxidation | *** | *** | ** | ** | *** | | | | |

$^{+}$ Abbreviation of environmental attributes: $R_h$, heterotrophic respiration; $R_t$, soil total respiration; $Q_{10}$, temperature sensitivity of heterotrophic respiration.

**Table S4.** The correlations between the structure of each functional gene group involved soil C decomposition and N cycling processes and each environmental attribute revealed by Mantel test. The functional community structure was determined by GeoChip hybridization. Significance is represented by *** when p < 0.01, ** when p <0.05 and * when p <0.10.

| | | GeoChip | | | Metagenome based EcoFUN-MAP | | | Metagenomic sequencing | | |
|---|---|---|---|---|---|---|---|---|---|---|
| | Attributes+ | $R_h$ | $R_t$ | $Q_{10}$ | $R_h$ | $R_t$ | $Q_{10}$ | $R_h$ | $R_t$ | $Q_{10}$ |
| | Carbon cycling | ** | ** | ** | | ** | | ** | ** | * |
| | Carbon degradation | ** | ** | ** | * | ** | | ** | * | * |
| | Starch | ** | ** | ** | ** | ** | | ** | * | |
| | Pectin | ** | ** | ** | ** | ** | ** | | | |
| | Hemicellulose | ** | ** | ** | * | ** | * | ** | ** | |
| **C degradation** | Cellulose | ** | ** | ** | | ** | ** | | | |
| | Chitin | ** | ** | ** | | ** | | | ** | |
| | Other | ** | ** | ** | | * | | * | ** | * |
| | Vanillin/lignin | ** | ** | ** | * | ** | | | | |
| | Ammonification | ** | ** | ** | ** | * | | ** | ** | *** |
| | Anammox | ** | | | | | | *** | *** | ** |
| | Assimilatory N reduction | ** | ** | ** | | ** | * | *** | *** | * |
| **N cycling** | Denitrification | ** | ** | ** | | | | | | |
| | Dissimilatory N reduction | ** | ** | ** | | * | | | | |
| | Nitrification | ** | ** | ** | | | | *** | *** | * |
| | Nitrogen fixation | ** | ** | ** | | | | | | ** |
| **P utilization** | P utilization | ** | ** | ** | | * | | *** | ** | |
| | Adenylylsulfate reductase | *** | *** | *** | | | | | | |
| **S metabolism** | Sulfur assimilation | | ** | *** | ** | | | | *** | |
| | Sulfite reduction | *** | *** | *** | | * | | | | *** |
| | Sulfide oxidation | *** | *** | *** | | | * | | | |

[+] Abbreviation of environmental attributes: $R_h$, heterotrophic respiration; $R_t$, soil total respiration; $Q_{10}$, temperature sensitivity of heterotrophic respiration.

**Table S5.** The enzyme/protein encoded by biogeochemical cycling genes shown in Figures. 2, S8 S9 and Tables S3, S4.

| Gene category | Subcategory | Gene name | Enzyme/protein encoded |
|---|---|---|---|
| | Glyoxylate cycle | *AceA* | Isocitrate lyase |
| | Glyoxylate cycle | *AceB* | Malate synthase A |
| | Starch | glucoamylase | Glucoamylase |
| | Starch | *cda* | Cyclomaltodextrinase |
| | Starch | *amyA* | Alpha-amylase |
| | Starch | *amyX* | Pullulanase |
| | Starch | *nplT* | Neopullulanase |
| | Starch | *apu* | Amylopullulanase |
| | Starch | isopullulanase | Isopullulanase |
| | Starch | *pula* | Pullulanase, extracellular |
| | Hemicellulose | xylanase | Xylanase |
| | Hemicellulose | mannanase | Beta-mannanase |
| | Hemicellulose | *xyla* | Xylose isomerase |
| | Hemicellulose | *ara* | Arabinofuranosidase |
| | Pectin | pectinase | Pectinase |
| | Pectin | pectin lyase | Pectin lyase |
| | Pectin | *Pg* | Polygalacturonase |
| | Pectin | *pel_Cdeg* | Pectin lyase |
| | Pectin | *rgh* | Rhamnogalacturonase |
| | Pectin | *pme* | Pectinesterase |
| | Pectin | exopolygalacturonase | Exopolygalacturonase |
| C degradation | Pectin | *RgaE* | Lipolytic enzyme |
| | Pectin | *rgl* | Polysaccharide lyase |
| | Pectin | pectate lyase | Pectate lyase |
| | Pectin | endopolygalacturonase | Endopolygalacturonase |
| | Cellulose | *axe* | Acetyl xylan esterase |
| | Cellulose | cellobiase | Cellobiase |
| | Cellulose | endoglucanase | Endoglucanase |
| | Cellulose | cellulase | Cellulase |
| | Cellulose | exoglucanase | Exoglucanase |
| | Camphor | camdcab | Camphor 5-monooxygenase |
| | Terpenes | *limeh* | Limonene-1,2-epoxide hydrolase |
| | Terpenes | *lmo* | Limonene 1,2-monooxygenase |
| | Terpenes | *cdh* | Carveol dehydrogenase |
| | Cutin | cutinase | Cutinase |
| | Chitin | acetylglucosaminidase | Acetylglucosaminidase |
| | Chitin | chitin deacetylase | Chitin deacetylase |
| | Chitin | chitinase | Chitinase |
| | Vanillin/Lignin | *vana* | Vanillate monooxygenase |
| | Vanillin/Lignin | *vdh* | Vanillin dehydrogenase |
| | Vanillin/Lignin | phenol oxidase | Phenol oxidase |
| | Vanillin/Lignin | ligninase | Ligninase |
| | Vanillin/Lignin | *glx* | Glyoxal oxidase |
| | Vanillin/Lignin | *mnp* | Manganese peroxidase |

| | | | |
|---|---|---|---|
| C fixation | Bacterial Microcompartments | *CsoS2* | Carboxysome |
| | Calvin cycle | *rubisco* | RuBisCo |
| | Calvin cycle | FBPase | Fructose-1 6-bisphosphatase |
| | Calvin cycle | *PRK* | Phosphoribulokinase |
| | Reductive acetyl CCoA | *codh* | Carbon monoxide dehydrogenase |
| | Reductive acetyl CCoA | *fthfs* | Tetrahydrofolate formylase |
| | Multiple systems | *pcc* | Propionyl-CoA carboxylase |
| | reductive tricarboxylic acid cycle | *aclb* | ATP citrate lyase |
| | reductive tricarboxylic acid cycle | *mdh* | Malate dehydrogenase |
| N cycling | Ammonification | *gdh* | Glutamate dehydrogenase |
| | Ammonification | *urec* | Urease |
| | Anammox | *hzsa* | Hydrazine synthase |
| | Anammox | *hzo* | Hydrazine oxidoreductase |
| | Assimilatory N reduction | *narb* | Nitrate reductase |
| | Assimilatory N reduction | *NiR* | Nitrite reductase |
| | Assimilatory N reduction | *nira* | Ferredoxin-nitrite reductase |
| | Assimilatory N reduction | *nirb* | Nitrite reductase |
| | Assimilatory N reduction | *nasa* | Assimilatory nitrate reductase |
| | Denitrification | *norb* | Nitric-oxide reductase |
| | Denitrification | *nirk* | Copper containing nitrite reductase |
| | Denitrification | *nirs* | Cytochrome cd1 nitrite reductase |
| | Denitrification | *cnorB* | Nitric oxide reductase |
| | Denitrification | *nosz* | Nitrous oxide reductase |
| | Denitrification | *narg* | Respiratory nitrate reductase |
| | Dissimilatory N reduction | *nrfa* | Ammonia-forming nitrate reductase |
| | Dissimilatory N reduction | *napa* | Periplasmic nitrate reductase |
| | Nitrification | *amoa* | Ammonia monooxygenase |
| | Nitrification | *hao* | Hydroxylamine oxidoreductase |
| | Nitrogen fixation | *nifh* | Dinitrogenase |
| P utilization | Phosphorus utilization | phytase | Phytase |
| | Phosphorus utilization | *ppx* | Exopolyphosphatase |
| | Phosphorus utilization | *ppk* | Polyphosphate kinase |
| S metabolism | Adenylylsulfate reductase | APS_AprA | Adenylylsulfate reductase |
| | Adenylylsulfate reductase | *AprA* | Adenylylsulfate reductase |
| | Adenylylsulfate reductase | APS_AprB | Adenylylsulfate reductase |
| | Sulfur assimilation | cysteine_synthase | Cysteine synthase |
| | Sulfur assimilation | ATP_sulphurylase | ATP sulphurylase |
| | Sulfur assimilation | PAPS_reductase | Phosphoadenosine phosphosulfate reductase |
| | Sulfite reduction | *cysI* | Sulfite reductase |
| | Sulfite reduction | *cysJ* | Sulfite reductase |
| | Sulfide oxidation | *sqr* | Sulfide-quinone reductase |
| | sulfite reduction | *dsrb* | Dissimilatory sulfite reductase |
| | sulfite reduction | *Sir* | Sulfite reductase |
| | sulfite reduction | *dsra* | Dissimilatory sulfite reductase |
| | Sulfur Oxidation | *sox* | Sulfur oxidation cycle enzymes |

**Table S6.** Soil carbon pools (state variables) in the MEND model.

| Soil carbon pool | Abbreviation | Variable name in governing equations |
|---|---|---|
| Particulate organic matter decomposed by oxidative enzymes | $POM_1$ | $P_1$ |
| Particulate organic matter decomposed by hydrolytic enzymes | $POM_2$ | $P_2$ |
| Mineral-associated organic matter | MOM | $M$ |
| Dissolved organic matter | DOM | $D$ |
| Active MOM interacting with DOM | QOM | $Q$ |
| Active microbial biomass | MBA | $BA$ |
| Dormant microbial biomass | MBD | $BD$ |
| Oxidative enzymes decomposing $POM_1$ | $EP_1$ | $EP_1$ |
| Hydrolytic enzymes decomposing $POM_2$ | $EP_2$ | $EP_2$ |
| Enzymes decomposing MOM | EM | $EM$ |

**Table S7.** Governing equations of each soil carbon pool in the MEND model

| Governing Equation | Eq# |
|---|---|
| $\frac{dP_1}{dt} = I_{P1} + (1-g_D) \cdot F_{12} - F_1$ | (S1) |
| $\frac{dP_2}{dt} = I_{P2} - F_2$ | (S2) |
| $\frac{dM}{dt} = (1-f_D) \cdot (F_1 + F_2) - F_3$ | (S3) |
| $\frac{dQ}{dt} = F_4 - F_5$ | (S4) |
| $\frac{dD}{dt} = I_D + f_D \cdot (F_1 + F_2) + g_D \cdot F_{12} + F_3 + (F_{14,EP1} + F_{14,EP2} + F_{14,EM}) - F_6 - (F_4 - F_5)$ | (S5) |
| $\frac{dBA}{dt} = F_6 - (F_7 - F_8) - (F_9 + F_{10}) - F_{12} - (F_{13,EP1} + F_{13,EP2} + F_{13,EM})$ | (S6) |
| $\frac{dBD}{dt} = (F_7 - F_8) - F_{11}$ | (S7) |
| $\frac{dEP_1}{dt} = F_{13,EP1} - F_{14,EP1}$ | (S8) |
| $\frac{dEP_2}{dt} = F_{13,EP2} - F_{14,EP2}$ | (S9) |
| $\frac{dEM}{dt} = F_{13,EM} - F_{14,EM}$ | (S10) |
| $\frac{dCO_2}{dt} = (F_9 + F_{10}) + F_{11}$ | (S11) |
| $\frac{d}{dt}(P_1 + P_2 + M + Q + D + BA + BD + EP_1 + EP_2 + EM) = I_{P1} + I_{P2} + I_D - (F_9 + F_{10} + F_{11})$ | (S12) |

The state variables (C pools) are described in Table S6; Eq. S11 indicates the total heterotrophic respiration flux and Eq. S12 expresses the overall mass balance of the system. The transformation fluxes are elucidated by Eqs. S13–S26 in Table S8.

**Table S8.** Component fluxes in the MEND model (parameters are described in Table S9)

| Flux description | Equation | Eq# |
|---|---|---|
| Particulate organic carbon (POC) pool 1 ($P_1$) decomposition ($F_1$) | $F_1 = \frac{Vd_{P1} \cdot EP_1 \cdot P_1}{K_{P1} + P_1}$ | (S13) |
| POC pool 2 ($P_2$) decomposition | $F_2 = \frac{Vd_{P2} \cdot EP_2 \cdot P_2}{K_{P2} + P_2}$ | (S14) |
| Mineral-associated organic carbon (MOC, $M$) decomposition | $F_3 = \frac{Vd_M \cdot EM \cdot M}{K_M + M}$ | (S15) |
| Adsorption ($F_4$) and desorption ($F_5$) between dissolved organic carbon (DOC, $D$) and adsorbed DOC (QOC, $Q$) | $F_4 = k_{ads} \cdot \left(1 - Q/Q_{\max}\right) \cdot D$<br>$F_5 = k_{des} \cdot (Q/Q_{\max})$ | (S16)<br>(S17) |
| DOC ($D$) uptake by microbes | $F_6 = \frac{1}{Y_g}\left(Vg + V_m\right)\frac{D \cdot BA}{K_D + D}$ | (S18) |
| Dormancy ($F_7$) and reactivation ($F_8$) between active (MBA) and dormant (MBD) microbial biomass ($BA$ and $BD$) | $F_7 = \left[1 - D/(K_D + D)\right] \cdot V_m \cdot BA$<br>$F_8 = D/(K_D + D) \cdot V_m \cdot BD$ | (S19)<br>(S20) |
| MBA ($BA$) growth respiration ($F_9$) and maintenance respiration ($F_{10}$) | $F_9 = \left(\frac{1}{Yg} - 1\right)\frac{V_g \cdot BA \cdot D}{K_D + D}$<br>$F_{10} = \left(\frac{1}{Yg} - 1\right)\frac{V_m \cdot BA \cdot D}{K_D + D}$ | (S21)<br>(S22) |
| MBD ($BD$) maintenance respiration | $F_{11} = \beta \cdot V_m \cdot BD$ | (S23) |
| MBA ($BA$) mortality | $F_{12} = \gamma \cdot V_m \cdot BA$ | (S24) |
| Synthesis of enzymes for $P_1$ ($EP_1$, $F_{13,\mathrm{EP1}}$), enzymes for $P_2$ ($EP_2$, $F_{13,\mathrm{EP2}}$), and enzymes for $M$ ($EM$, $F_{13,\mathrm{EM}}$) | $F_{13,EP1} = P_1/(P_1 + P_2) \cdot p_{EP} \cdot V_m \cdot BA$<br>$F_{13,EP2} = P_2/(P_1 + P_2) \cdot p_{EP} \cdot V_m \cdot BA$<br>$F_{13,EM} = p_{EM} \cdot V_m \cdot BA$ | (S25) |
| Turnover of enzymes ($EP_1$, $EP_2$, $EM$) | $F_{14,EP1} = r_E \cdot EP_1$<br>$F_{14,EP2} = r_E \cdot EP_2$<br>$F_{14,EM} = r_E \cdot EM$ | (S26) |

Notes: Italic symbols like $F_i$ represent component fluxes in equations. Italic symbols $P_1$, $P_2$, $M$, $Q$, $D$, $BA$, $BD$, $EP_1$, $EP_2$, and $EM$ are state variables (soil carbon pools, see Table S6) in equations.

**Table S9 Microbial-ENzyme Decomposition (MEND) model parameters**

| ID | Parameter | Description | Units | Eq# |
|---|---|---|---|---|
| 1 | $LF_0$ | Initial fraction of $P_1$, $LF_0 = P_1/(P_1+P_2)$ | — | |
| 2 | $r_0$ | Initial active fraction of microbes, $r_0 = BA/(BA+BD)$ | — | |
| 3 | $fINP$ | Scaling factor for litter input rate | — | |
| 4 | $Vd_{P1}$ | Maximum specific decomposition rate for $P_1$ | mg C mg$^{-1}$ C h$^{-1}$ | S13 |
| 5 | $Vd_{P2}$ | Maximum specific decomposition rate for $P_2$ | mg C mg$^{-1}$ C h$^{-1}$ | S14 |
| 6 | $Vd_M$ | Maximum specific decomposition rate for $M$ | mg C mg$^{-1}$ C h$^{-1}$ | S15 |
| 7 | $K_{P1}$ | Half-saturation constant for $P_1$ decomposition | mg C cm$^{-3}$ soil | S13 |
| 8 | $K_{P2}$ | Half-saturation constant for $P_2$ decomposition | mg C cm$^{-3}$ soil | S14 |
| 9 | $K_M$ | Half-saturation constant for $M$ decomposition | mg C cm$^{-3}$ soil | S15 |
| 10 | $Q_{\max}$ | Maximum sorption capacity | mg C cm$^{-3}$ soil | S16 |
| 11 | $K_{ba}$ | Binding affinity, Sorption rate $k_{ads} = k_{des} \times K_{ba}$ | (mg C cm$^{-3}$ soil)$^{-1}$ | S16 |
| 12 | $k_{des}$ | Desorption rate | mg C cm$^{-3}$ soil h$^{-1}$ | S17 |
| 13 | $r_E$ | Turnover rate of $EP_1$, $EP_2$, and $EM$ | mg C mg$^{-1}$ C h$^{-1}$ | S26 |
| 14 | $p_{EP}$ | $[V_m \times p_{EP}]$ is the production rate of $EP$ ($EP_1 + EP_2$), $V_m$ is the specific maintenance rate for $BA$ | — | S25 |
| 15 | $fp_{EM}$ | $fp_{EM} = p_{EM}/p_{EP}$ , $[V_{mt} \times p_{EM}]$ is the production rate of $EM$ | — | S25 |
| 16 | $f_D$ | Fraction of decomposed $P_1$ and $P_2$ allocated to $D$ | — | S3 |
| 17 | $g_D$ | Fraction of dead $BA$ allocated to $D$ | — | S1 |
| 18 | $V_g$ | Maximum specific uptake rate of $D$ for growth | mg C mg$^{-1}$ C h$^{-1}$ | S21 |
| 19 | $\alpha$ | $= V_m / (V_g + V_m)$ | — | S22 |
| 20 | $K_D$ | Half-saturation constant for microbial uptake of $D$ | mg C cm$^{-3}$ soil | S18 |
| 21 | $Y_g(T_{\text{ref}})$ | Intrinsic carbon use efficiency at reference temperature ($T_{\text{ref}}$) | — | S28 |
| 22 | $k_{Yg}$ | Slope for $Y_g$ dependence of temperature | 1/°C | S28 |

| | | | | |
|---|---|---|---|---|
| 23 | $Q_{10}$ | $Q_{10}$ for temperature response function | — | S28 |
| 24 | $\gamma$ | Max microbial mortality rate = $V_m \times \gamma$ | — | S24 |
| 25 | $\beta$ | Ratio of dormant maintenance rate to $V_m$ | — | S23 |
| 26 | $\psi_{A2D}$ | Soil water potential (SWP) threshold for microbial dormancy; both $\psi_{A2D}$ & $\psi_{D2A} < 0$ | −MPa | S30 |
| 27 | $\tau$ | $\psi_{D2A} = \psi_{A2D} \times \tau$, $\psi_{D2A}$ is the SWP threshold for microbial resuscitation | — | S30 |
| 28 | $\omega$ | Exponential in SWP function for microbial dormancy or resuscitation | — | S30 |

Notes: The column "Eq#" lists the major equation # (see Table S7 and S8) in which each parameter is used.

**Table S10.** Objective functions used for different response variables in the MEND model parameterization.

| Response Variable | Description of Response Variable | Objective Function for Each Response Variable | |
|---|---|---|---|
| | | Control | Warming |
| $R_h$ | Heterotrophic Respiration | $R^2$ between Simulated $R_h$ and Observed $R_h$ | $R^2$ between Simulated $R_h$ and Observed $R_h$ |
| MBC | Microbial Biomass Carbon | MARE < 20%<br>MBC_mean = 0.025 mg C $cm^{-3}$<br>(MBC = 2% SOC)<br>MBC_mean_simulated = 0.02 | MARE < 5%<br>MBC_mean = 0.02*0.84 = 0.017 mg C $cm^{-3}$ |
| EnzCo | Oxidative Enzyme Concentration (EnzC) | Correlation (r) between Simulated EnzC and Observed gene abundance (DNA concentration × relative abundance) | MARE between Simulated EnzC and Expected EnzC<br>Expected EnzC = Simulated EnzC at Control × RR |
| EnzCh | Hydrolytic Enzyme Concentration | Correlation (r) between Simulated EnzC and Observed gene abundance | MARE between Simulated EnzC and Expected EnzC |

Notes: RR is the response ratio of gene abundance under warming to that under control. $R^2$ denotes the coefficient of determination, MARE is the mean absolute relative error, see Methods Eqs. 3–4.
